# Mechanics of Esophageal Retraction During Anterior Cervical Discectomy and Fusion

**DOI:** 10.64898/2026.04.23.720008

**Authors:** Chihtong Lee, Alex R. Flores, Mert Çulcu, Alexander E. Ropper, Raudel Avila

## Abstract

Dysphagia, difficulty swallowing due to irritation or damage to the esophagus, is one of the most common complications following anterior cervical discectomy and fusion (ACDF), the most frequently performed cervical spine procedure in the United States. Surgical retraction hardware imposes sustained compression on the esophagus during surgery, generating nonuniform stress and strain fields that may contribute to temporary postoperative soft tissue damage. Current intraoperative assessment relies on visual inspection and manual inspection by the surgical team and does not provide quantitative measures of esophageal deformation, strain, or retraction displacement. Here, we present a comprehensive mechanics analysis of esophageal compression during ACDF that integrates experiments on esophageal phantoms, nonlinear finite element modeling, and theoretical thick-wall scaling relationships. Modeling results quantify peak contact pressures and corresponding stress distributions, identifying conditions under which circumferential strain in the compressed esophageal wall increases sharply as localized pressures approach the upper physiological range (∼6-17 kPa). Parametric investigation of retractor blade width, placement depth, and polymeric biocompatible coating properties demonstrates that targeted, yet mechanically simple, design modifications can help to attenuate strain concentrations. In particular, the introduction of compliant polymeric coatings redistributes contact loads and reduces peak wall stress by up to 20% relative to unbuffered blades (17 kPa to 13.5 kPa). Increasing blade width from 20 mm to 50 mm further decreases peak interface stress from 2.48 kPa to 0.45 kPa, corresponding to an 82% reduction. Reducing these stresses may help limit mechanically induced complications such as postoperative dysphagia. Experiments performed on esophageal phantoms with embedded pressure sensors replicate surgical ACDF retraction protocols under displacement-controlled conditions. This setup establishes physiologically relevant loading and enables quantitative validation of computational predictions by correlating measured voltage output with contact pressure and esophageal deformation. Measured relationships between applied retraction displacement, contact pressure, and tissue deformation govern stress amplification during ACDF retraction. Together, these results establish a predictive mechanics framework that links retractor blade design variables to esophageal stress fields, providing quantitative criteria to mitigate soft tissue damage during ACDF.

**HIGHLIGHTS:**

- 2D and 3D finite element models quantify esophageal wall stress during anterior cervical discectomy and fusion (ACDF) retraction.
- Retractor blade geometry influences stress distribution, with wider blades reducing localized tissue loading by up to 82% likely associated with post-surgical dysphagia.
- Compliant polymeric buffer layers attenuate pressure and smoothen stress gradients to reduce peak tissue loading by up to 20% during retraction.

## 1. INTRODUCTION

Anterior cervical discectomy and fusion (ACDF) is a well-established surgical procedure for treating herniated or degenerated intervertebral discs in the cervical spine. Degeneration and collapse of herniated discs located between adjacent vertebrae (C1 through C7) produces sustained neural compression and chronic radicular pain (Figure 1a). ACDF alleviates these symptoms through removal of damaged discs and insertion of a structural bone or synthetic grafts, accessed through an anterior cervical approach that minimizes disruption and damage to posterior musculature and neural structures (Cleveland Clinic, 2024). Between 2010 and 2022, approximately 1.2 million cervical spine surgeries were performed in the contiguous United States, of which 61.2% were ACDF procedures (743,300 cases) (Ibrahim et al., 2025). As the population ages and the incidence of cervical spine trauma increases, surgical demand is expected to rise (Neifert et al., 2020). Although ACDF provides effective symptom relief, postoperative complications remain common, and the projected growth in surgical volume underscores the need to better understand and mitigate mechanically induced complications and reduce recovery times.

**Figure 1:**
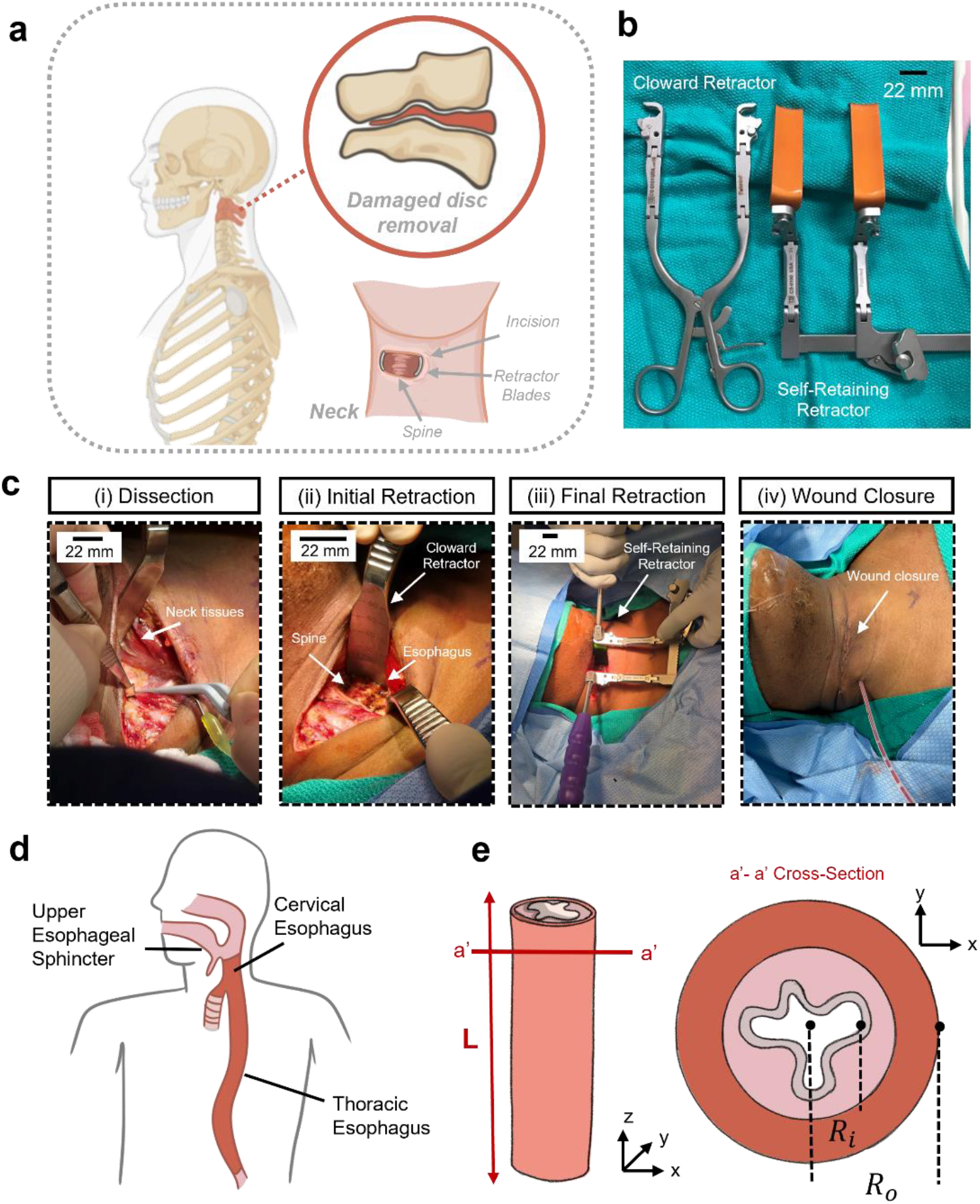
Retraction mechanism of neck tissues in anterior cervical discectomy and fusion (ACDF). (a) Anatomical location of the cervical spine and schematic illustration of intervertebral disc removal and replacement during ACDF. (b) Representative surgical retractors used to maintain the operative field of view during the ACDF procedure, including two examples of self-retraction instruments. (c) Optical images of the surgical workflow illustrating key stages of the procedure, including initial dissection, tissue retraction, discectomy, and post-surgical wound closure. (d) Illustration of the anatomical position of the cervical esophagus within the neck. (e) Cross-sectional schematic of the cervical esophagus represented as a simplified layered structure used for the mechanical modeling framework.

The anterior approach in ACDF requires lateral retraction of cervical soft tissues, including the trachea and, most critically, the esophagus, to expose the cervical spine. Rigid retractor blades, commonly fabricated from stainless steel or medical-grade titanium (Thompson Retractor) and mounted on self-retaining instruments (Figure 1b), are inserted through the incision and laterally displace the soft tissues. Typical retraction displacements range from 15 to 20 mm, establishing a surgical field of view that provides unobstructed visualization and clearance for instrument access to the intervertebral discs. In clinical practice, retraction aims to optimize surgical exposure to access the spine, while the mechanical response and consequences of the compressed esophageal wall remain largely unquantified. The sequential stages of dissection, initial retraction, and sustained holding during ACDF for cervical spondylotic myelopathy (Figure 1ci–1ciii), illustrated using intraoperative pictures from a patient treated at an academic research hospital, generate localized contact pressures at the interface between the rigid retractor blade and the soft esophageal wall, where excessive deformation or tissue collapse can persist beyond wound closure (Figure 1civ). ACDF surgeries often involve prolonged retraction, with reported mean operative durations of 159 ± 24.3 min (n = 39) for single-level ACDF and 220 ± 33.4 min (n = 12) for two-level ACDF (Chung et al., 2018). These extended retraction periods can generate nonuniform stress and strain fields within the esophageal wall, which may contribute to tissue discomfort, inflammation, and postoperative edema.

Postoperative difficulty swallowing, or dysphagia, is among the most frequently reported complications following ACDF. Reported overall morbidity rates range from 13.2% to 19.3% (Vaishnav et al., 2019; Tsalimas et al., 2022). Within these complications, dysphagia occurs in 1.7% to 9.5% of procedures, with incidence as high as 21.3% at 2-year follow-up in some cohorts (Riley et al., 2005). Based on national surgical ACDF procedures, these rates correspond to thousands (2,000–27,000) of affected patients annually in the United States. Despite its prevalence, dysphagia is often regarded as an inherent risk of anterior cervical exposure (Vaishnav et al., 2019; Riley et al., 2005; Tsalimas et al., 2002). However, mechanics consequences between retraction/compression and postoperative esophageal dysfunction remain poorly quantified. In particular, the relationships between retraction displacement, contact pressure, and stress and strain development in the esophageal wall have not been systematically resolved.

Prior studies have examined biomechanical and clinical contributors to postoperative dysphagia following ACDF. Several investigations identify retractor geometry and placement as factors influencing esophageal deformation and potential tissue injury. Papavero et al. (2007) evaluated direct pressure transmission during retraction and its association with postoperative swallowing disturbances. Although the overall correlation was low, the authors suggested that dysphagia may arise from highly localized mechanical loading rather than global tissue compression. Additional clinical studies highlight additional risk factors including: cervical graft (plate) design, resulting esophageal adhesion to the hardware and cervical spine during surgery, and patient demographics (Lee et al., 2005; Rhyne et al., 2007; Nguyen et al., 2022).

Recent studies highlight the role of surgical hardware–tissue interactions, as well as the duration and magnitude of retraction, in contributing to soft tissue trauma and inflammation (Fogel and McDonnell, 2005; Perez-Roman et al., 2021; Liu et al., 2017). In a retrospective analysis of 37 ACDF patients, Ramos-Zúñiga et al. (2015) reported that manual retraction techniques using protective blade systems were associated with lower rates of soft tissue complications, suggesting that controlled modulation of mechanical loading influences the esophageal response. These findings indicate that retraction mechanics introduces modifiable variables in surgical practice. However, quantitative relationships between these design parameters and resulting stress and strain fields in the esophageal wall remain insufficiently characterized.

We test the hypothesis that ACDF-induced retraction generates measurable mechanical stress signals within the esophageal wall, and that these signals, originating from localized contact pressure, stress amplification, and sustained deformation, can be quantitatively resolved using nonlinear finite element modeling to interpret the mechanical consequences of surgical procedures and hardware choices. Rather than viewing retraction as a uniform esophageal displacement, we examine how spatially localized compression governs adverse tissue loading under continuous deformation.

The mechanical behavior of the esophagus presents a nontrivial modeling problem due to its nonlinear, multilayered, and anisotropic soft tissue fibrous structure, with substantial interpatient variability (Durcan et al., 2022; Yang et al., 2006; Sommer et al., 2013; Ren et al., 2021). The cervical esophagus extends from the upper esophageal sphincter to the thoracic segment (Figure 1d) and can be idealized in cross-section as a thick-walled cylindrical tube with inner radius *R*_*i*_ and outer radius *R*_*o*_ (Figure 1e). Structurally, it consists of a mucosa– submucosa composite layer and an outer muscular layer, each exhibiting distinct mechanical characteristics approximated by the phenomenological hyperelastic models. The mucosa–submucosa has a higher collagen content and exhibits different stiffness and deformation behavior than the surrounding muscle (Stavropoulou et al., 2012; Sommer et al., 2013). Although finite element (FE) methods enable multiscale modeling of nonlinear, incompressible constitutive behavior and layered esophageal geometry, direct *in vivo* quantification of esophageal strain and contact pressure during ACDF retraction in the operating room (Figure 1c) remains limited by hardware access constraints and bio-integrated electronic sensor capabilities. To address this limitation and complement the computational analysis, we develop a soft-tissue physical phantom that reproduces the essential geometry and bulk mechanical properties of the cervical esophagus and neck. This experimental platform enables controlled measurement of retraction displacement and contact pressure fields and provides validation of computational predictions.

In this study, we integrate clinical insight with computational biomechanical analysis and experimental validation to advance the understanding of esophageal retraction during ACDF. The manuscript is organized as follows. The Methods section describes (1) the development and analysis of finite element models of the esophagus, (2) the selection and characterization of hyperelastic constitutive models, and (3) experimental approaches designed to capture the complex mechanics of surgical retraction in a controlled system. The Results section presents key findings from the in-silico simulations and corresponding experimental measurements, including validation of pressure distributions along the esophageal wall. To further interpret these findings, we develop an analytical thick-walled cylinder approximation for a hyperelastic tube under pressure, providing additional insight into stress gradients across the esophageal wall during retraction. Collectively, these results establish a quantitative framework for evaluating surgical design parameters-including blade width, retraction displacement, and polymer coating thickness-that govern localized strain within the esophageal wall. Ultimately, we will provide mechanistic insight into how retractor design will influence esophageal tissue loading, with the potential to reduce postsurgical complications such as dysphagia from the hardware standpoint.

## 2. METHODS

### 2.1 Geometric Esophageal Model and Boundary Conditions

The cervical esophagus, located between the upper esophageal sphincter and the thoracic esophagus, is modeled as a thick-walled cylindrical tube of length *L* = 60 mm, consistent with reported anatomical dimensions (Durcan et al., 2022).

Figure 2a shows a schematic of the observed deformation of the cervical esophagus to yield appropriate surgical field of view, along the length of the 60 mm section. The undeformed configuration is defined by an inner radius *R*_*i*_ = 5 mm, corresponding to a luminal diameter of 10 mm (Ren et al., 2021; Sommer et al., 2013), and an outer radius *R*_0_ = *R*_*i*_ + *t*_*m*_ + *t*_*mus*_ where *t*_*m*_ = 1.2 mm denotes the thickness of the mucosa–submucosa layer and *t*_*mus*_ = 2.3 mm denotes the thickness of the muscularis layer (Sommer et al., 2013; Yang et al., 2006 (a)). The total wall thickness is 3.5 mm. The esophagus is modeled as a two-layer deformable solid in the reference configuration as

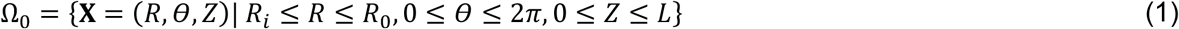

**Figure 2:**
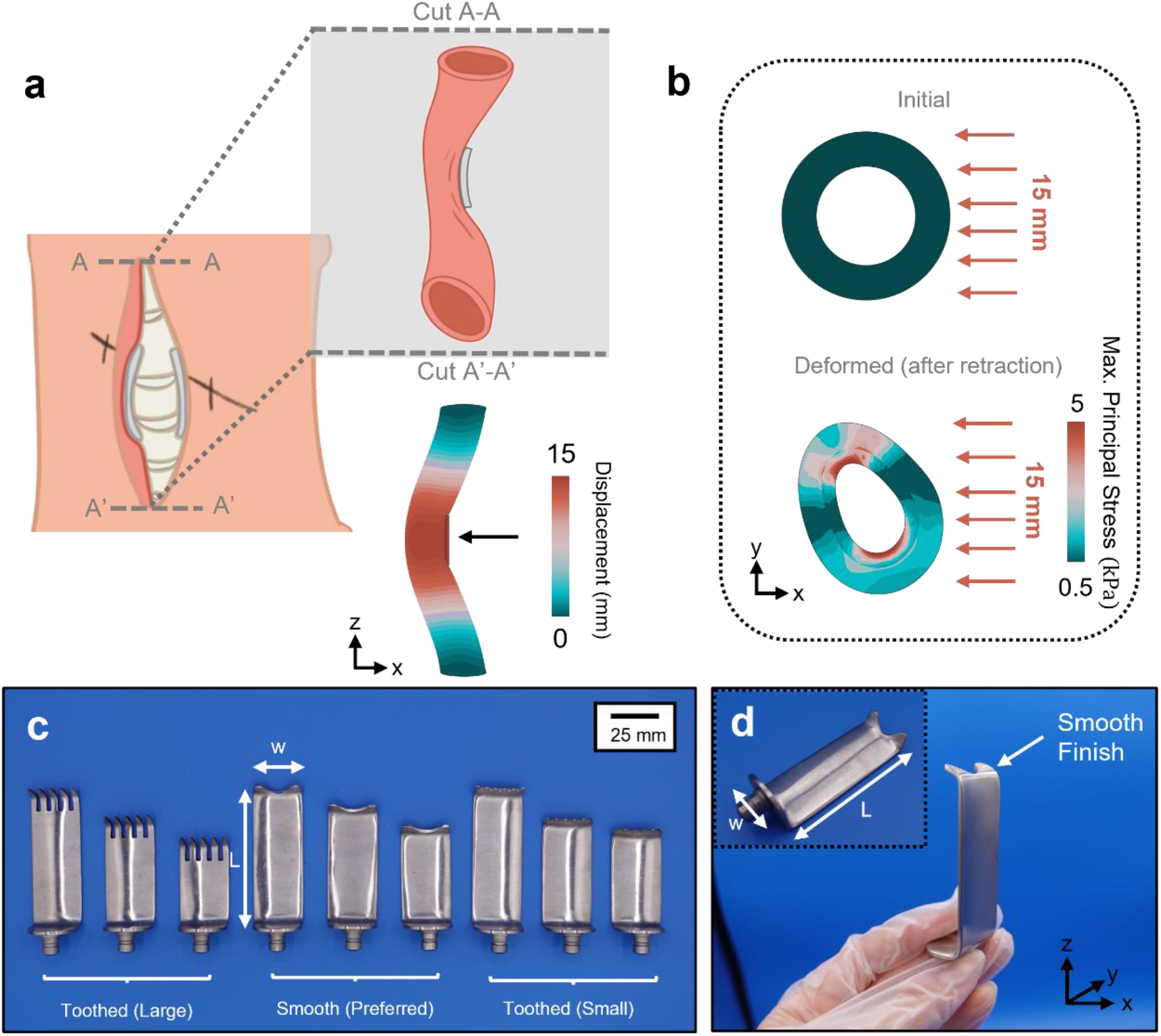
Esophageal deformation and retractor blade variations. (a) Schematic illustration of esophageal deformation resulting from surgical opening and interaction with the retractor blade. (b) Cross-sectional view of the esophagus in the finite element model showing the initial configuration (top) and the maximum principal stress in the deformed configuration (bottom) following 15 mm of surgical retraction. (c) Representative variations in retractor blade design and geometry used clinically, with blade length and width differing by up to 50 mm. (d) Optical image of the 20 mm smooth-finish retractor blade used in the retraction experiments.

The motion is defined by the deformation mapping as

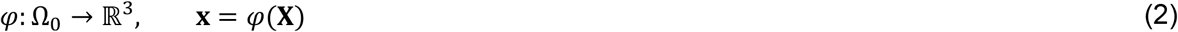

with displacement field **u**(**X**) = *φ*(**X**) − **X** and the deformation gradient **F** = ∇**x***φ*. The Green–Lagrange strain tensor follows

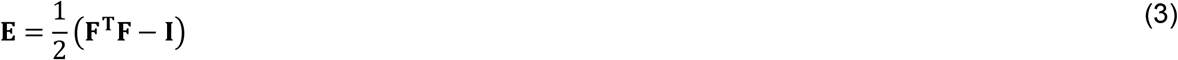

and the Cauchy stress tensor ***σ*** is calculated from the hyperelastic strain energy function (refer to the material model section) and used to evaluate stress, contact pressure, and strain concentrations induced by blade-driven compression (Figure 2a-b). To represent tissue fixation at the upper esophageal sphincter and the thoracic segment, Dirichlet boundary conditions are prescribed at the proximal and distal ends

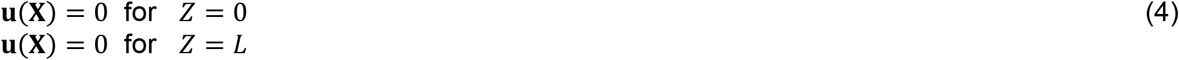

These constraints suppress rigid-body motion and enforce physical anchoring of the cervical segment. The retractor blade occupies a rigid surface Γ_*b*_ positioned laterally at the mid-span of the esophageal segment, centered at *Z* = *L*/2. T Figure 2c-d shows the three types of retractor blades used in surgery, with the most desirable blade applied to the lateral side being the smooth blade. The blade in FE models is idealized as a rigid rectangular prism with a prescribed width *w* and length *L*_*b*_. The blade undergoes displacement-controlled lateral retraction. Its rigid-body motion is defined by

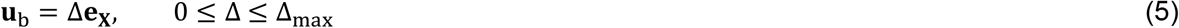

Where **e**_**X**_ is the unit vector in the lateral direction and Δ_max_= 15 mm represents the imposed surgical retraction displacement. All other rigid-body motion components are suppressed, such that the blade kinematics are fully determined by the scalar parameter Δ. Contact between the rigid blade surface Γ_*b*_ and the deformable esophageal outer surface Γ_*c*_ ⊂ *∂*Ω is modeled in Abaqus/Standard using surface-to-surface contact with finite sliding. Normal contact behavior follows a unilateral (non-penetration) constraint consistent with hard contact, represented in continuum form by the Kuhn–Tucker conditions

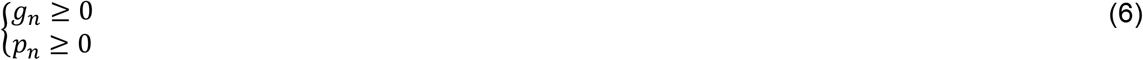

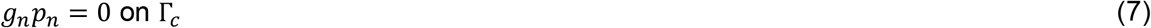

where *g*_*n*_(**x**) is the normal gap function (positive in separation) and *p*_*n*_(**x**) is the normal contact pressure. Tangential behavior is prescribed as frictionless, such that tangential traction *t*_*t*_ vanishes and the contact traction acting on the esophageal surface reduces to

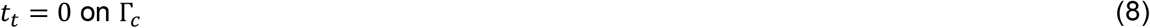

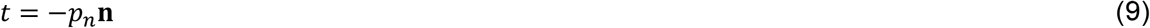

where **n** denotes the outward unit normal to the esophageal surface in the current configuration.

### 2.2 Finite Element Modeling

Two-dimensional (2D) and three-dimensional (3D) finite element simulations were performed in Abaqus/Standard (Dassault Systemes, Simulia) to resolve large deformation contact mechanics between the rigid retractor blade and the deformable esophageal wall. In both cases, the esophagus was modeled as a two-layered hyperelastic solid with a thick-walled cylindrical geometry (Supplementary Figure 1a). The rigid retractor blade and optional polymeric coating layer (Ecoflex 00-30) were included in the assembly to evaluate the sensitivity of geometric and material parameters during ACDF retraction. The 2D models consisted of approximately 30,000 bilinear eight-node hybrid plane strain elements (CPE8H). The retractor blade was modeled as a rigid surface, while an optional Ecoflex layer was discretized using hybrid elements in soft buffer thickness studies. In studies with the buffer, nonlinear geometry (NLGEOM) was enabled to capture evolving contact conditions and changing geometries. Self-contact was defined along the luminal surface to prevent nonphysical penetration of the esophageal wall under large deformation. Retraction was imposed via displacement of the rigid blade, while the bottom and back face of the tissue were constrained only to parallel motion to its axis. To evaluate radial and circumferential stress distributions, uniform external pressures of 5, 10, and 20 kPa were applied to the outer surface of the esophageal wall. These loading conditions approximate the compressive pressures generated during surgical retraction and enable analytical evaluation of stress development across the wall thickness.

In the 3D models, only the esophagus was modeled as a deformable body to resolve stress distribution during retraction (Supplementary Figure 1b). The deformable domains were discretized using eight-node linear brick hybrid elements (C3D8H), appropriate for nearly incompressible hyperelastic materials. A hybrid formulation was employed to mitigate volumetric locking under large strain and near-incompressible response. The model contained approximately 18,000 elements, with local mesh refinement applied in the anticipated contact region. Contact between the rigid blade and the esophageal outer surface was defined using surface-to-surface finite-sliding interaction with frictionless tangential behavior. Normal contact behavior followed a hard-contact formulation. Retraction was imposed via lateral displacement control of the rigid blade in Eq. 5, with quasi-static solution assumptions used to enforce equilibrium at each increment. Nonlinear geometric effects were accounted for using large-deformation kinematics. Convergence was monitored by the residual norms of force and displacement using the default Newton–Raphson iterative scheme.

### 2.3 Ecoflex 00-30 Esophageal Phantom Fabrication

An experimental platform was developed to replicate displacement-controlled retraction of the cervical esophagus and to measure contact pressure during blade compression. The esophagus was fabricated as a hollow cylindrical tube with a length *L* = 100 mm, inner radius *R*_*i*_ = 5 mm, and wall thickness *t* = 3.5 mm, consistent with anatomical dimensions used in the computational models. The phantom was fabricated using a custom 3D-printed polylactic acid (PLA) mold coated with a release agent and allowed to dry for 30 min prior to casting. The esophageal phantom was cast using Ecoflex 00-30 silicone elastomer (Smooth-On), selected for its nonlinear, nearly incompressible hyperelastic response representative of idealized soft biological tissue. The elastomer components (8.8 oz, Part A and Part B, in a 1:1 weight ratio) were mixed for 3 min to ensure homogeneity. Red pigment (Smooth-On SilcPig) was added to improve optical contrast during deformation tracking. The mixture was poured into the mold and cured at room temperature for 4 h prior to demolding (Figure 3a), following well-established fabrication protocols and curing conditions (Yang et al., 2013). Following fabrication, the cylindrical phantom was embedded within a bulk Ecoflex matrix to represent the surrounding cervical neck tissue. The matrix was cast within a 3D-printed PLA enclosure with dimensions 50 × 120 × 50 mm (Figure 3b). The esophageal tube was positioned 20 mm below the superior surface and laterally offset by 10 mm to approximate the anatomical relationship between the esophagus and the surgical approach (Figure 3c-3d). After curing, a 40 mm-deep incision was performed along the superior surface to allow controlled blade insertion and resection during ACDF simulated retraction testing (Figure 3d).

**Figure 3:**
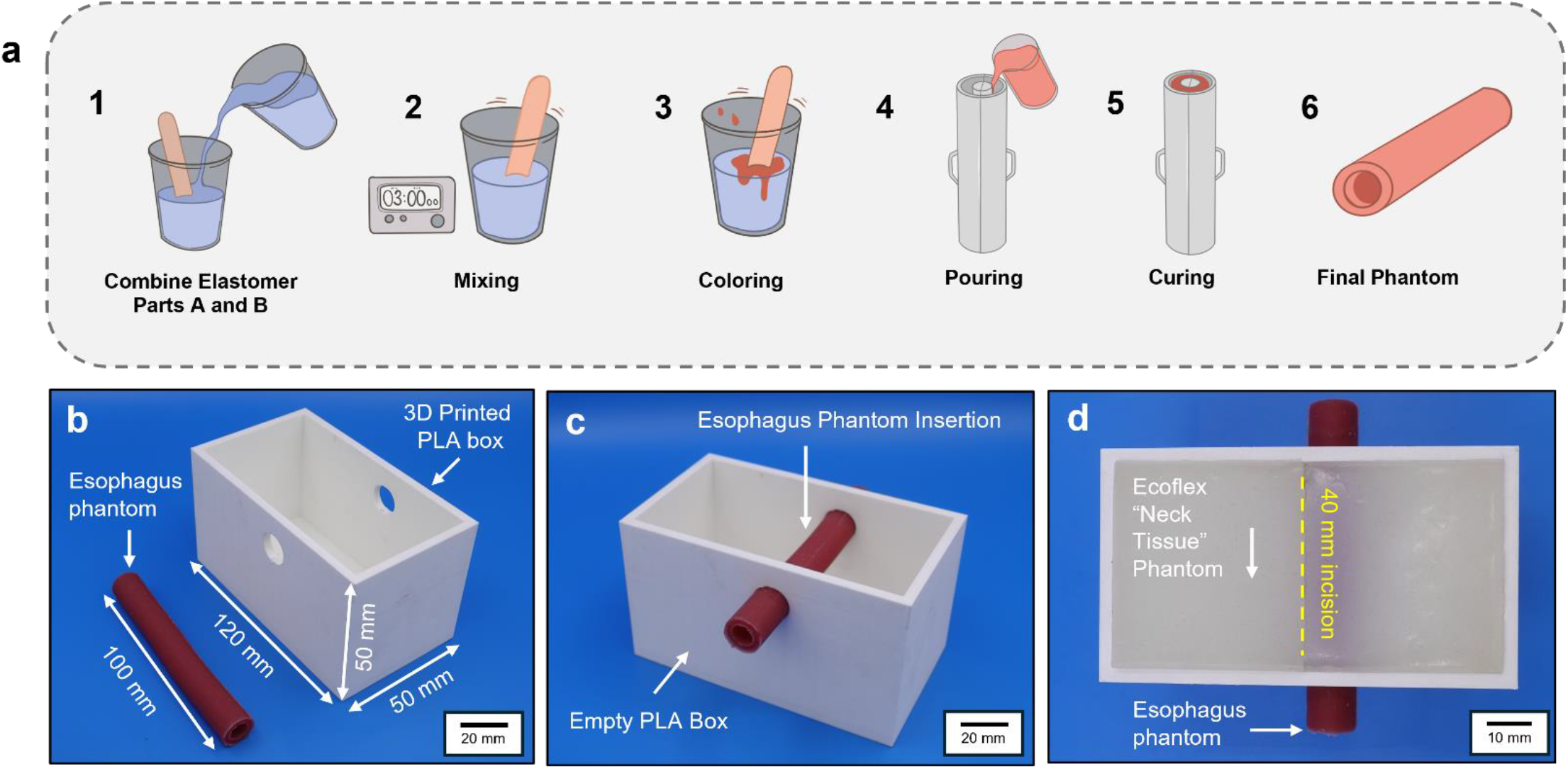
Esophageal phantom fabrication. (a) Schematic illustration of the fabrication process for the esophageal phantom. (b) Final phantom components integrated within a 3D-printed PLA enclosure. (c) Esophageal phantom positioned inside the PLA box for retraction testing. (d) Final phantom configuration with neck tissue and a 40 mm deep incision (yellow dashed line) used to replicate the surgical access pathway.

### 2.4 Pressure Sensor Positioning and Calibration

Thin-film pressure sensors (Model: DF9-16; Weight = 0.04 g; Diameter = 10 mm, Length = 19 mm) were conformably attached to the medial surface of the esophageal phantom to measure normal contact pressure induced by the rigid retractor blade. Figure 4a illustrates the data acquisition setup used to measure contact pressures during retraction. Insets show the thin-film pressure sensor positioned next to a U.S. quarter coin for scale, as well as the full phantom retraction setup indicating the incision line and sensor placement relative to the esophageal model. Retraction was imposed under displacement-controlled loading consistent with the computational boundary conditions in Eq. 5. Sensor leads were soldered using a fine-tip soldering iron to minimize thermal damage and preserve electrical contacts. The unloaded resistance of the force-sensitive resistor (FSR) was measured as 0.968 MΩ using a digital multimeter. The sensor was connected to a voltage divider circuit consisting of a fixed series resistor, *R*_1_ = 10 kΩ, and the variable sensor resistance, *R*_FSR_. The circuit was powered using the NI ELVIS data acquisition system (Figure 4a). Output voltage was recorded in LabVIEW 2025 at a sampling rate of 20 Hz over a variable acquisition window (10–60 seconds). The measured output voltage followed the voltage divider relation

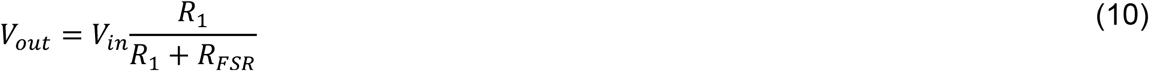

where *V*_in_ is the supplied input voltage and *R*_FSR_ decreases with increasing applied normal force. Under this configuration, *V*_out_ increases monotonically with applied load. Calibration was performed by applying known masses of 100 g, 200 g, 400 g, 800 g, and 1000 g sequentially to the active sensing region (diameter: 10 mm) (Figure 4b). Each mass was applied in three independent trials. To ensure uniform load distribution across the sensing surface area (*A* = 78 mm^2^), the sensor was mounted onto a rigid 3D-printed PLA fixture and secured using medical-grade silicone tape (Solventum 2477) to prevent slip and off-axis loading. Applied mass *m* was converted to force using *F* = *mg* where *g* = 9.81 m · s^−2^. Contact pressure was then computed as

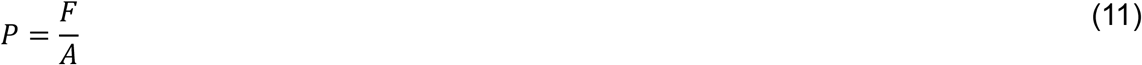

**Figure 4:**
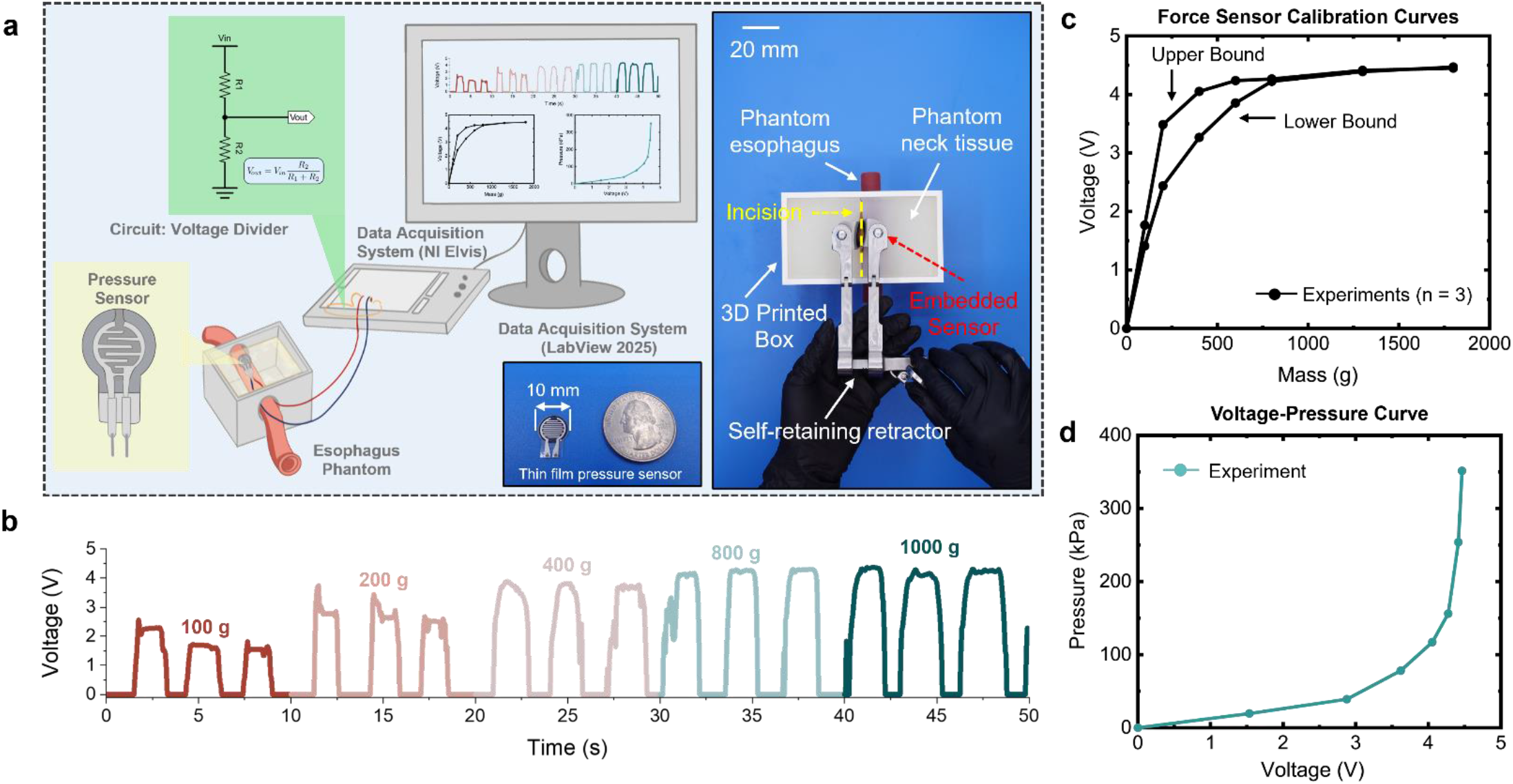
Experimental setup for ADCF-like retraction in phantom esophagus. (a) Schematic of the sensor circuitry and experimental setup used to quantify pressure during retraction. Inset: optical image of the thin-film pressure sensor shown alongside a U.S. quarter for scale. (b) Temporal voltage response of the pressure sensor under applied loads of 100, 200, 400, 800, and 1000 g, demonstrating the sensor response during incremental loading. (c) Calibration curve relating applied loads to the measured sensor output showing the upper and lower bounds. (d) Converted voltage–pressure relationship curve obtained from the calibration procedure.

The resulting voltage–pressure relationship was obtained by correlating *V*_out_ with calculated pressure values (Figure 4c–d). Pressure values were obtained from the measured sensor voltage using the calibration curve derived from the voltage–load experiments. The relationship between output voltage and pressure was approximated using an exponential best-fit function of the form

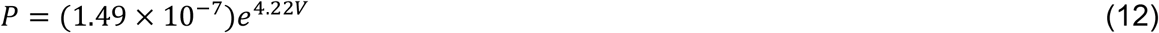

The upper and lower bounds of the calibration curve were used to define the uncertainty range for subsequent pressure measurements during retraction experiments.

### 2.5 Mechanical Characterization of Ecoflex 00-30

Uniaxial tensile tests were performed to characterize the nonlinear hyperelastic response of Ecoflex 00-30 used in the phantom model. Testing was conducted using an ARES G-2 rheometer equipped with a 20 N load cell under ambient laboratory conditions (22 °C, 54% relative humidity). Rectangular specimens (30 × 3 × 1 mm) were fabricated from cured Ecoflex 00-30 sheets and tested in uniaxial tension (Figure 5a-b). A laser-cut alignment template was used to ensure consistent gauge length and minimize geometric variability between specimens. The effective gauge length was 10 mm. Three samples were subjected to monotonic tensile loading up to 500% engineering strain at a constant strain rate of 1 mm/s with force and displacement data recorded at 30 Hz (Figure 5c-d). To reduce stress concentrations at the grips and minimize slippage, paper support frames were bonded to the specimen ends prior to testing and removed after mounting. This configuration ensured uniform axial loading within the gauge region. The engineering strain, stretch, and engineering stress were calculated as

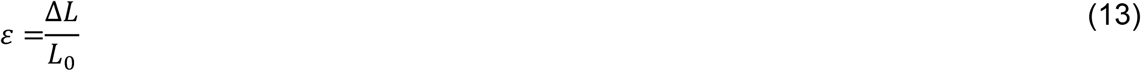

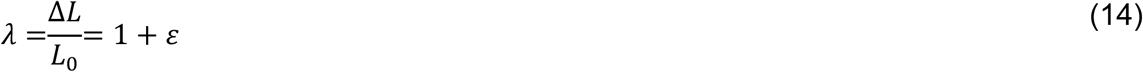

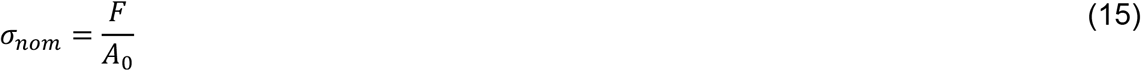

**Figure 5:**
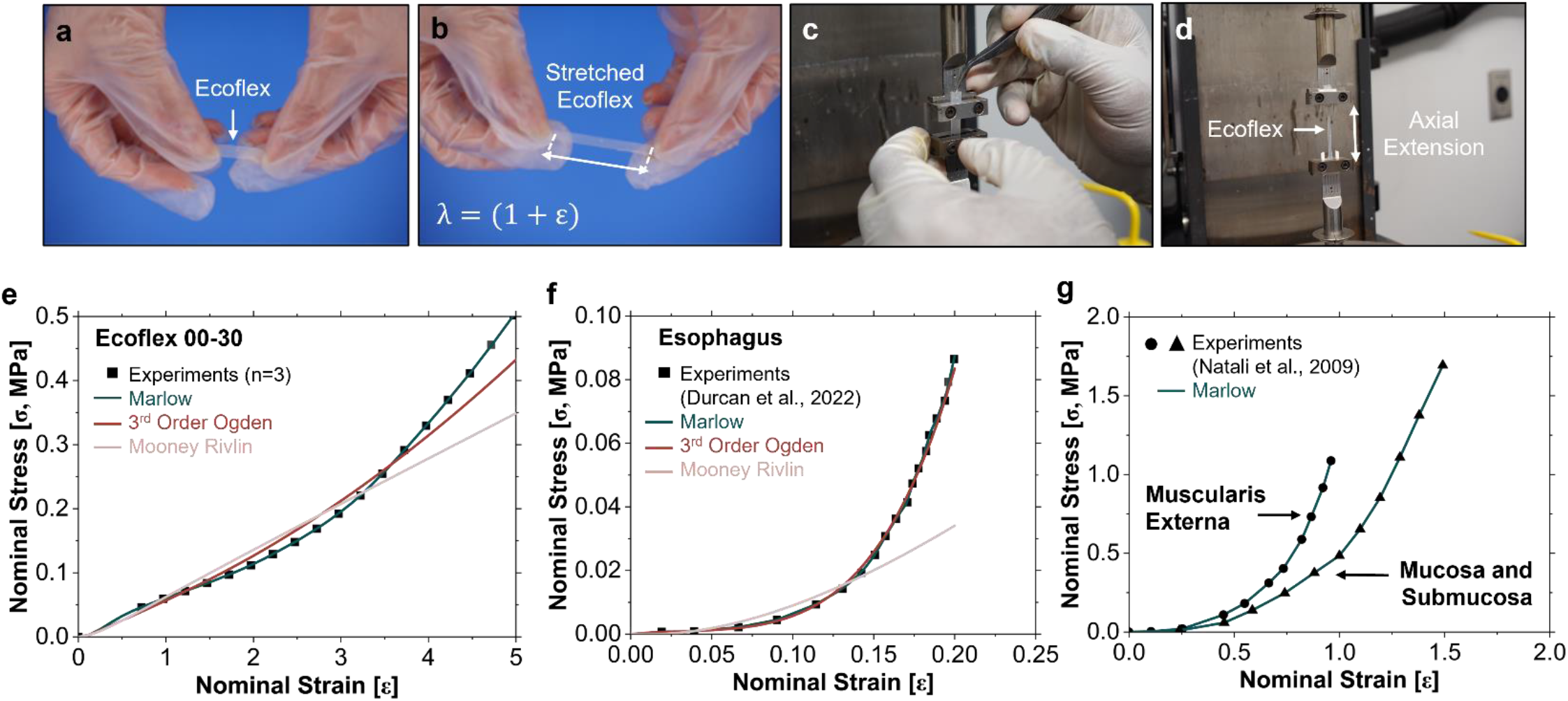
Experimental calibration and constitutive model fitting. (a) Ecoflex 00-30 specimens (30 mm gauge length) prepared for mechanical characterization. (b) Ecoflex 00-30 specimen under manual uniaxial stretch. (c) Uniaxial tensile testing setup using a rheometer (ARES G2) equipped with a 20 N load cell. (d) Close-up view of the tensile testing configuration used for soft material characterization. (e) Uniaxial tensile stress– strain response of Ecoflex 00-30 (n = 3) with hyperelastic constitutive model fits using Marlow, Ogden (N = 3), and Mooney–Rivlin strain energy functions. (f) Nominal stress–strain response of esophageal tissue reported by Durcan et al. (2022) with the corresponding hyperelastic constitutive model fit. (g) Nominal stress–strain response of esophageal tissue from tensile testing data reported by Natali et al. (2009) with the corresponding Marlow constitutive model fit.

Where *L*_0_ is the initial gauge length, *F* is the measured force, and *A*_0_ is the initial cross-sectional area. For constitutive material calibration in ABAQUS, the nominal stress–stress response was used as input to define the hyperelastic material parameters. Representative stress–strain curves for Ecoflex 00-30 are shown in Figure 5e, comparing experimental data with three distinct hyperelastic strain energy functions (i.e., Marlow, Ogden, and Mooney Rivlin).

### 2.6 Hyperelastic Material Models

#### Esophagus

The esophageal wall is modeled as a nonlinear hyperelastic material undergoing finite deformation. The constitutive response is defined in terms of a strain-energy density function *W*(**F**) where **F** is the deformation gradient and *J* = det(**F**). To account for volumetric–isochoric decoupling under large deformation, the strain energy function is expressed as

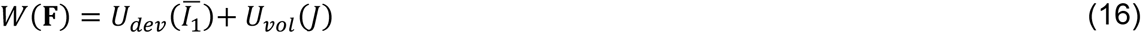

where 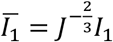 is the first deviatoric invariant of the modified right Cauchy–Green tensor and *I*_1_ = tr(**C**) with **C** = **F**^**T**^**F**. The volumetric contribution is governed by *J*, ensuring near incompressibility of the soft tissue. The esophageal material properties are approximated using the Marlow hyperelastic formulation implemented in Abaqus/Standard. In this model, the deviatoric strain-energy function 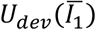 is constructed directly from experimental uniaxial stress–strain data without assuming a predefined functional form. Uniaxial tensile data reported by Durcan et al. (2022) were used to define the deviatoric response (Figure 5f), with an excellent fit to the Ogden and Marlow hyperelastic models. The volumetric component is defined through the Poisson ratio *ν* = 0.49, enforcing near-incompressible behavior consistent with soft biological tissues. The volumetric strain-energy contribution takes the quadratic form

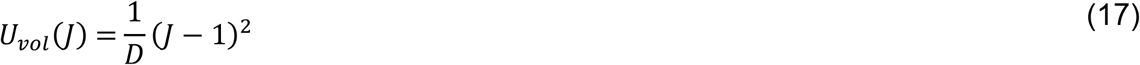

where *D* is related to the bulk modulus. The Marlow formulation was selected because it preserves the experimentally observed stress–strain response without requiring parameter fitting to a prescribed polynomial or exponential representation. This approach reduces extrapolation error at large strains while maintaining numerical stability in finite-deformation contact simulations. In the initial two-dimensional model, bilayer formulation was used to visualize esophageal behavior under compression. Uniaxial tensile data reported by Natali et al. (2009) was used for the subsequent deviatoric response (Figure 5g) for the muscularis external and the mucosa and submucosa layers.

#### Ecoflex 00-30

For completeness, conventional parametric hyperelastic models were evaluated to describe the nonlinear, nearly incompressible behavior of Ecoflex. The compressible Ogden model of order *N* = 3 is defined as

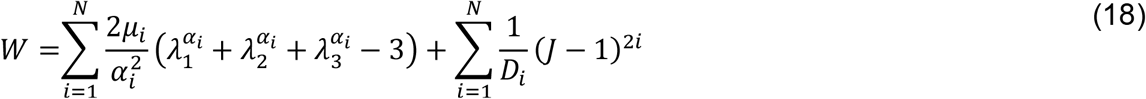

where *λ*_*i*_ are the principal stretches, and *μ*_*i*_, *α*_*i*_, and *D*_*i*_ are material parameters determined through nonlinear regression. The two-parameter Mooney-Rivlin model is given by

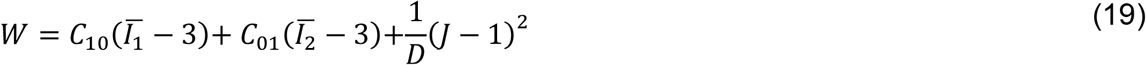

where 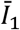 and 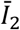 are the modified first and second invariants of **C**, and *C*_10_, *C*_01_, *D*_1_ are material constants. Although the Ogden and Mooney–Rivlin formulations reproduce the uniaxial stress–strain response over limited strain intervals (Figure 5e-f) and are retained for closed-form analytical derivations of thick-walled cylinder stresses, the data-driven Marlow hyperelastic model was adopted for all subsequent numerical simulations of Ecoflex 00-30. All material constants are provided in Tables 1 and 2.

**Table 1.** Material constants of Ecoflex 00-30.

Ecoflex Material Properties from Experimental Data
| Ogden (N=3) | $\mu_1$[MPa] | $\alpha_1$ | $\mu_2$[MPa] | $\alpha_2$ | $\mu_3$[MPa] | $\alpha_3$ |
| --- | --- | --- | --- | --- | --- | --- |
|  | 1.017 | -0.390 | 0.004 | 4.006 | -1.020 | -0.790 |
| Mooney-Rivlin | $C_{10}$[MPa] | | $C_{01}$[MPa] | | $D_1$[MPa<sup>-1</sup>] | |
|  | 0.0349 |  | -0.034 |  | 0 |  |

**Table 2.**
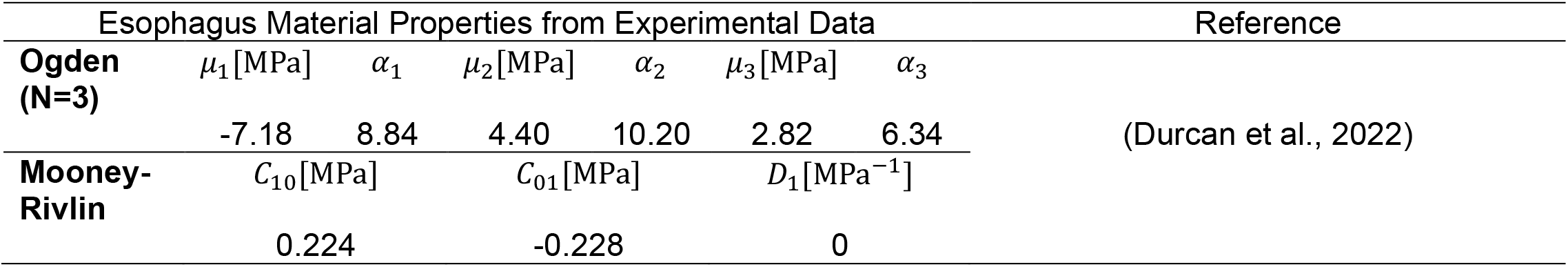
Material constants of Esophageal Tissue (longitudinal)

## 3. RESULTS

### 3.1 Parametric Studies – Modeling Surgical Hardware

A parametric analysis was conducted to quantify the stresses of the esophageal wall under displacement-controlled lateral retraction by a rigid surgical blade. A prescribed lateral displacement of Δ = 15 mm, applied normal to the longitudinal axis of the esophageal segment, was used to represent clinically relevant retraction magnitudes during ACDF surgery (Figure 6a).

**Figure 6:**
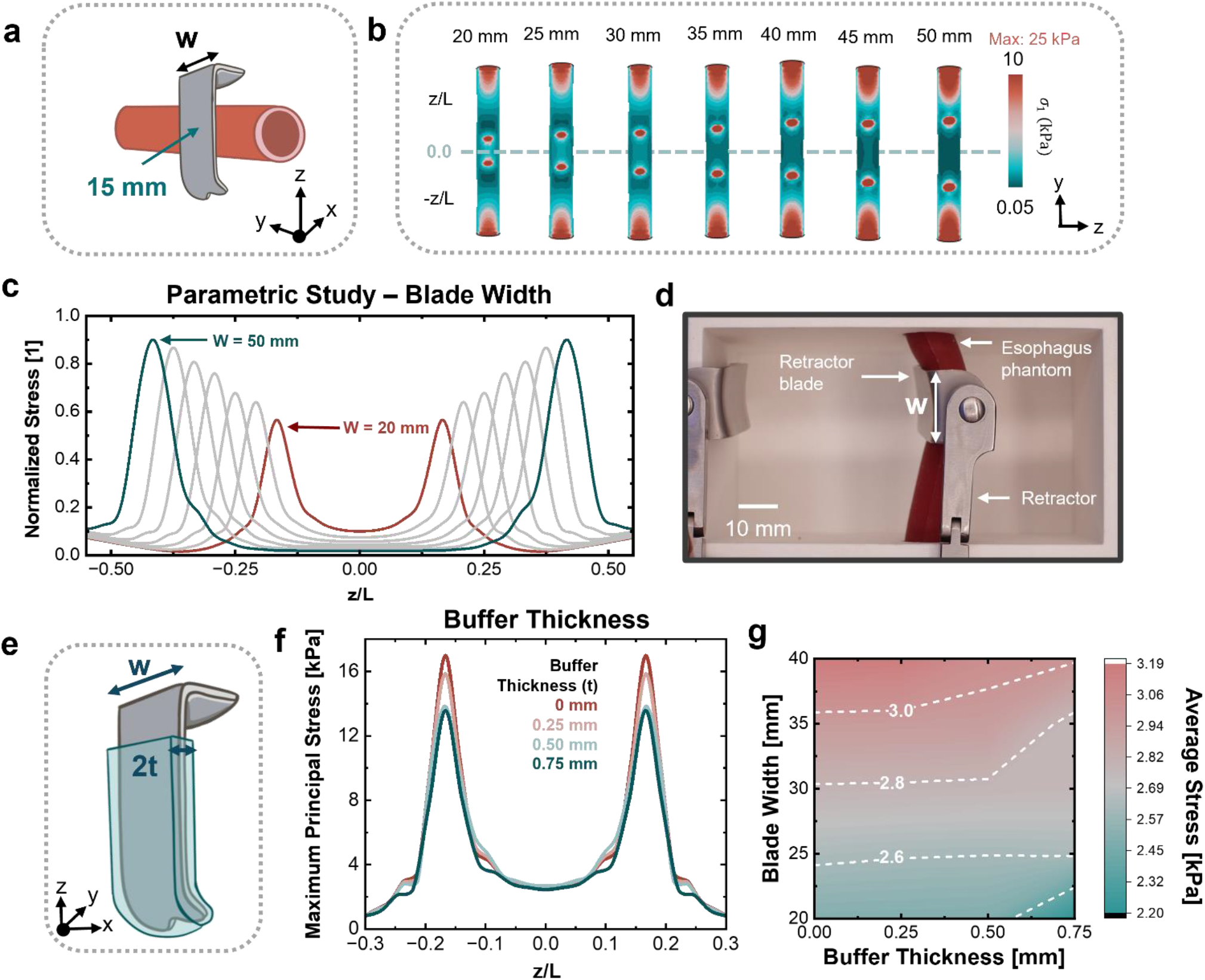
Finite element analysis of retractor blade geometry and soft buffer sleeve. (a) Schematic illustration of a rigid retractor blade in contact with the esophagus. (b) Parametric evolution of stress fields in the esophageal wall with increasing blade width (left to right), ranging from 20 to 50 mm. (c) Normalized stress distribution along the normalized esophageal length obtained from the finite element simulations, showing two characteristic peaks corresponding to the contact regions where stresses are highest. (d) Experimental retraction of the esophagus qualitatively illustrating the resulting tissue displacement in the absence of surrounding neck tissues. (e) Schematic illustration introducing a polymeric buffer layer (or sleeve) between the blade and esophagus to reduce stress concentrations. (f) Maximum principal stress along the normalized esophageal length for three representative buffer layer thicknesses. (g) Parametric investigation of blade width and buffer thickness showing the bilinear surface plot of the average stresses in the esophagus.

#### 3.1.1 Blade Width and Localized Stress Response

Figure 6b shows the distribution of maximum principal stress along the outer surface of the esophageal wall under blade-induced retraction. The stress field is evaluated along the axial direction of the cylindrical esophageal segment to quantify spatial variations in tissue loading arising from the retractor–tissue contact as blade width increases from 20 mm to 50 mm. For all blade geometries, the stress distribution exhibits a pronounced local maximum beneath the central contact region, where the imposed displacement is transmitted directly into the soft esophageal wall. The elevated stress concentrations (10 kPa >) quantified near the superior and inferior edges of the esophageal segment arise from the fixed boundary conditions imposed at the axial ends of the model. To quantitatively compare blade geometries, stress profiles are presented in normalized form as a function of the nondimensional axial coordinate *z*/*L*, where *z* denotes the axial position along the esophagus and *L* is the total esophageal length (Figure 6c). Stresses were normalized by the maximum stress obtained along the esophageal surface under the reference loading condition of 25 kPa.

At the blade centerline 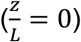, the maximum principal stress reaches 2.48 kPa for the narrow blade configuration (*w* = 20 mm). Increasing the blade width to 50 mm reduces the stress to 0.45 kPa, corresponding to an approximately 82% reduction. This decrease reflects the larger contact interface, which distributes the imposed displacement over a greater tissue area and thereby lowers localized stress concentrations. Figure 6c reveals two characteristic stress peaks for each blade geometry located near the blade edges, where the esophageal wall bends and bulges around the rigid contact interface. These edge regions generate localized stress concentrations as the imposed displacement is accommodated by bending and stretching of the surrounding tissue. For the widest blade (*w* = 50 mm), the edge stresses reach approximately (≈22.5 kPa), whereas for the narrower blade (*w* = 20 mm) the peak edge stresses are approximately (≈13.8 kPa). This behavior highlights an inherent geometric trade-off: increasing blade width reduces the peak stress beneath the blade center by distributing the load across a larger contact area but simultaneously shifts higher stresses toward the blade edges where the esophageal wall undergoes greater curvature and deformation. Although wider blades require greater total force to achieve the prescribed displacement, the region of the esophagus directly beneath the blade experiences lower stress levels.

The associated curved deformation of the esophageal wall under these loading conditions is illustrated experimentally in Figure 6d using a silicone esophageal phantom and a 20 mm blade. For visualization purposes, the surrounding tissue matrix was removed to isolate the deformation of the esophageal structure during ACDF retraction. The observed deformation patterns qualitatively agree with the displacement fields predicted by the finite element simulations, with the largest inward displacement occurring directly beneath the retractor blade and outward bulging developing along the blade edges. Determining whether clinical outcomes are governed primarily by peak localized stresses or by the spatially averaged stress distribution along the esophageal wall lies beyond the scope of the present study. In practice, postoperative dysphagia likely results from a combination of mechanical loading, tissue inflammation, surgical duration, and patient-specific anatomical factors. Nevertheless, the model provides a quantitative framework for evaluating readily adjustable surgical design parameters, such as retractor blade width, influence stress concentrations and deformation patterns in the esophageal wall.

#### 3.1.2 Blades with a Soft Polymeric Sleeve

Figure 6e presents a schematic illustrating the introduction of a compliant polymeric (Ecoflex 00-30) buffer sleeve layer applied to the surface of the surgical blade to reduce stress transfer to the esophageal tissue during retraction. The buffer is modeled as an incompressible soft elastomeric layer of thickness 2*t* conformally attached to the blade surface, thereby introducing an intermediate interface between the rigid surgical blade and the soft esophageal wall. This design choice reduces the stiffness mismatch at the blade–tissue interface and is intended to mitigate stress concentrations from bulging generated during ACDF retraction.

For a blade width of 20 mm, the influence of a compliant polymeric sleeve with thicknesses of 0.25 mm, 0.50 mm, and 0.75 mm was evaluated. The resulting stress fields show a moderate reduction in localized tissue loading at the blade–tissue interface. Specifically, the buffer sleeve design decreases the peak maximum principal stress by approximately 20%, from 17 kPa to 13.5 kPa, relative to the unbuffered configuration. Quantitatively, the 0.75 mm buffer layer reduces the peak stress by approximately 3.41 kPa. Although the overall shape of the stress profiles in Figure 6f remains similar, the buffered design attenuates stress concentrations near the blade edges, indicating that the compliant layer redistributes the applied displacement over a broader contact region.

To assess whether the effectiveness of the compliant buffer depends on blade geometry, additional simulations were performed across a two-dimensional parameter space varying blade widths while incrementally increasing buffer thickness. Figure 6g summarizes the combined influence of blade width and compliant buffer thickness on the average stress within the esophageal wall. The contour map indicates that average stress increases with blade width, reflecting the larger portion of tissue retracted by the contact as the blade spans a greater axial length. In contrast, increasing the buffer thickness consistently reduces the average stress by introducing a compliant interface that mitigates stiffness mismatch and redistributes the applied displacement across the tissue surface. The contours further reveal a bi-linear interaction between blade width and buffer thickness, most pronounced beyond a buffer thickness of approximately 0.5 mm. While wider blades reduce stresses within the central contact region (Figure 6c), they increase the spatially averaged stress transmitted to the esophageal wall due to the stress peaks that develop near the blade edges. The addition of a soft buffer layer partially offsets this effect by reducing the average stress across the contact region. These results highlight a mechanical trade-off between minimizing localized stress concentrations beneath the blade and limiting the overall stress transmitted to the surrounding tissue during retraction.

### 3.2 Experimental Retraction Studies with Esophageal Phantom

#### 3.2.1 Retraction Sequence with Self-Retaining Retractor

Experimental voltage measurements were performed using a self-retaining surgical retractor equipped with smooth blades positioned on the lateral and contralateral sides of the esophageal phantom. A five-stage stepwise opening protocol was implemented to characterize the pressure response during progressive lateral retraction (Figure 7a). Displacement increments were controlled by applying point markings along the retractor handle at 5 mm intervals, providing a visual reference for repeatable opening. The retractor was actuated over a total duration of approximately 10 s, with each displacement increment held for ∼2 s to allow the pressure sensor signal to stabilize before proceeding to the next stage. This loading protocol ensured that the measurements correspond to quasi-static deformation, thereby minimizing rate-dependent effects associated with rapid tissue deformation.

**Figure 7:**
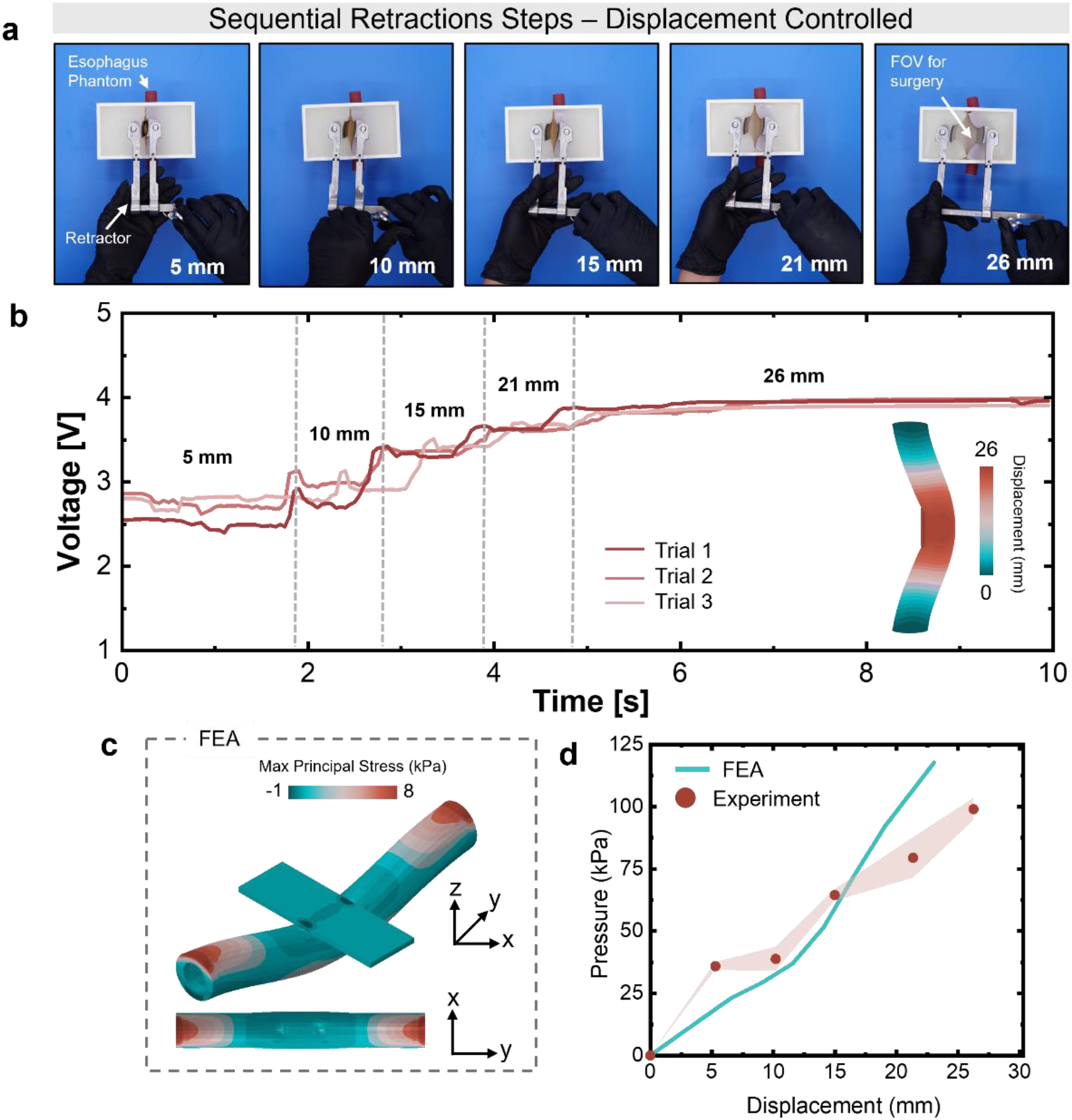
Experimental response of the ACDF phantom esophageal model. (a) Representative continuous retraction sequence using a self-retaining retractor applied to the esophageal phantom to achieve an opening of 26 mm. (b) Temporal sensor voltage response corresponding to discrete step openings during retraction from 5 to 26 mm. (c) Three-dimensional finite element model showing the maximum principal stress distribution in the esophagus subjected to 20 mm compressive deformation during simulated surgical opening. (d) Comparison of pressure–displacement curves obtained from finite element simulations and experimental measurements.

Figure 7b shows the measured voltage response of the pressure sensor during each stage of lateral retraction, corresponding to blade displacements ranging from 5 mm to 26 mm measured from the medial line of the surgical incision. For comparison, the displacement field predicted by the finite element model is shown as a contour. As the retraction displacement increases, the measured voltage rises monotonically, from 2.5 V to 4V, indicating an increase in contact pressure transmitted to the sensor. Initial baseline voltage drifts between 2.5 V and 2.8 V at 5 mm are attributed to variations in the initial placement of the retractor blade on the esophageal phantom. These trends follow the same monotonic voltage–pressure relationship established in the calibration curve described in Section 2.4, confirming that the sensor response remains consistent across the tested loading range. Three measurements demonstrate that progressive blade displacement produces increasing compressive loading at the esophagus interface, which is reflected by the corresponding increase in sensor output.

To enable direct comparison with the experimental measurements, a finite element model uses the Marlow hyperelastic constitutive model (Figure 5e) and boundary conditions consistent with the experimental configuration (Figure 7c) and can quantify the maximum principal stresses on the esophagus wall. The model represents the retractor blade as a rigid surface undergoing displacement-controlled lateral motion, while the esophageal phantom behaves as a deformable solid with fixed boundary conditions at the proximal and distal ends. Figure 7d compares the experimentally measured pressures with the contact pressures predicted by the finite element simulations across all displacement stages. Good agreement is obtained for blade displacements up to approximately 15 mm, indicating that the calibrated constitutive model captures the mechanical response of the phantom under moderate deformation. In this range, both experiment and simulation show a consistent increase in contact pressure with increasing blade displacement. At larger displacements, the simulations predict average contact pressures up to 37% higher than those measured experimentally. This discrepancy likely arises from geometric nonlinearities in the deformation of the phantom and surrounding neck tissue (Figure 7a), variations in sensor positioning at large displacements, and contact mechanics effects that are not captured by the rigid blade idealization and become increasingly significant at higher compression levels. Despite these deviations, the overall pressure–displacement relationship remains consistent between experiment and simulation, supporting the validity to extract pressure metrics that describe the mechanics of esophageal retraction.

#### 3.2.2 Continuous Retraction Study with a Soft Polymeric Sleeve

The influence of a compliant polymeric buffer layer applied to the rigid surgical blade was investigated experimentally to evaluate its ability to reduce contact pressures at the interface. These experiments were designed to cross-validate the trends predicted by the finite element simulations, which indicated that increasing the thickness of a compliant interfacial layer reduces localized stress concentrations beneath the blade. In the experiments, the buffer layer, an Ecoflex 00-30 elastomeric sheet, is bonded to the blade surface, introducing a thin but deformable intermediate layer between the rigid blade and the compliant esophageal phantom. Mechanically, this thin layer modifies the contact interface by increasing the effective separation between the rigid blade and the esophageal surface prior to retraction. Upon initial contact, the compliant layer undergoes elastic compression, generating a preload/prestress within the contact interface and within the pressure sensing system. Figure 8a shows an optical image of the proposed compliant sleeve design applied to the surgical blade, featuring a sleeve thickness of 1 mm.

**Figure 8:**
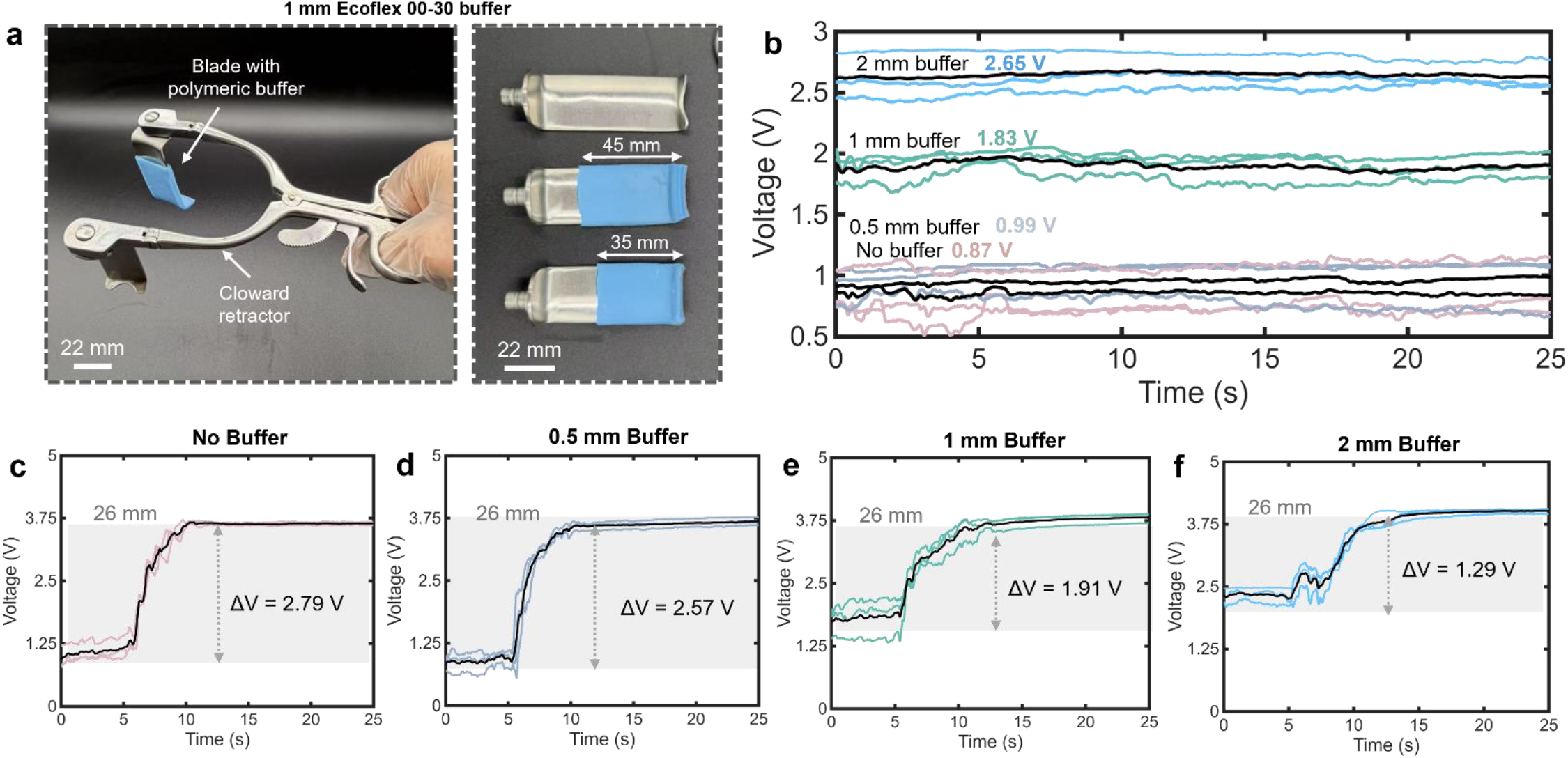
Experimental voltage response with buffered surgical blades. (a) Optical images of surgical blades featuring a 1 mm thick polymeric buffer sleeve mounted on a Cloward retractor; the sleeve can be designed to cover different lengths of the blade. (b) Baseline voltage measurements obtained with the blade inserted into the esophageal phantom for buffer layer thicknesses ranging from 0 to 2 mm. (c) Continuous opening voltage response during retraction without a buffer layer. (d) Continuous opening voltage response with a 0.5 mm buffer layer. (e) Continuous opening voltage response with a 1 mm buffer layer. (f) Continuous opening voltage response with a 2 mm buffer layer.

This effect is reflected in the baseline voltage response of the thin-film pressure sensor prior to blade displacement. Thicker buffer layers produce higher baseline voltage levels due to greater compressive deformation of the elastomer during initial contact. Figure 8b shows the baseline voltage levels measured for the four buffer conditions investigated: no buffer, 0.5 mm, 1 mm, and 2 mm Ecoflex layers. The average baseline voltages over three independent trials were 0.87 V, 0.99 V, 1.83 V, and 2.65 V, respectively, reflecting the increasing elastic precompression introduced by thicker polymeric layers.

ACDF-simulated retraction experiments were conducted over a 25 s time window to quantify the voltage response as a function of buffer layer thickness. Figures 8c–f show the temporal voltage response during retraction, exhibiting a monotonic increase over the first 10 s as the blade is displaced to 26 mm. After approximately 10 s, the signal reaches a plateau as the displacement is held constant and the contact pressure stabilizes. Each of the four conditions includes three independent trials in which the blade was returned to the initial position prior to repeating the retraction experiment. The averaged response for each condition is shown as the black curve, demonstrating excellent repeatability across trials.

Table 3 summarizes the total change in voltage measured between the initial blade insertion (baseline) and the fully retracted configuration (change) at 26 mm displacement for each buffer thickness. The addition of a 2 mm compliant buffer layer reduces the overall voltage increase by approximately 54% relative to the rigid blade alone. This reduction in signal amplitude indicates that a smaller portion of the imposed displacement is transmitted directly to the central blade–tissue contact region, consistent with the compliant layer redistributing the applied load over a larger effective contact area.

**Table 3.** Voltage changes in continuous retraction with introduction of polymeric buffer.

| | Baseline Voltage (V) | Change in Voltage ($\Delta V$) |
| --- | --- | --- |
| No buffer (rigid blade) | 0.87 | 2.79 |
| 0.5 mm | 0.99 | 2.57 |
| 1 mm | 1.83 | 1.91 |
| 2 mm | 2.65 | 1.29 |

The continuous voltage signals cannot be converted directly to absolute pressure values because the large retraction displacement generates non-uniform contact pressure distributions which influence the magnitude of the sensor response. Nevertheless, the relative scaling with respect to each baseline and the systematic decrease in signal amplitude with increasing buffer thickness indicate that a compliant buffering sleeve reduces the magnitude of interface loading during retraction. This behavior agrees with the computational predictions, which show that a soft layer reduces the interfacial stiffness mismatch between the rigid blade and the esophageal tissue. This stiffness redistribution smooths spatial stress gradients along the esophageal surface.

### 3.3 Thick Wall Semi-Analytical Approximation of Esophageal Compression

Consider a thick-walled circular esophagus with undeformed inner and outer radii *R*_*i*_ and *R*_*o*_, respectively (Figure 9a). Under a uniform external pressure *P*, the deformed radii are denoted by *r*_*i*_ and *r*_*o*_. The analysis assumes axisymmetric finite deformation and plane strain, such that *λ*_*z*_ = 1.

**Figure 9:**
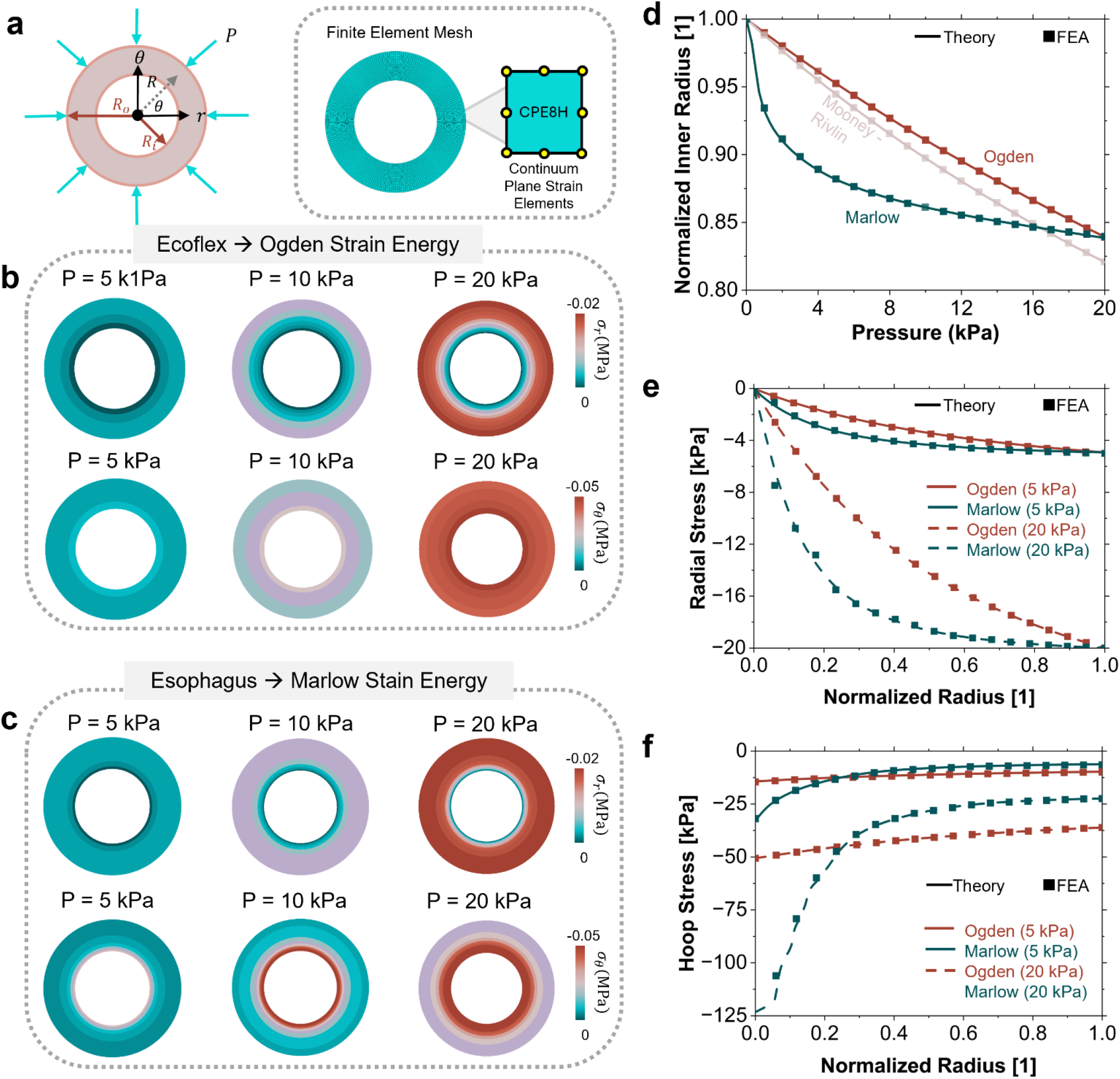
Semi-analytical theory for thick wall approximation. (a) Axisymmetric thick-walled cylinder subjected to external pressure *P*together with the corresponding finite element mesh used in the simulations. (b) Radial and hoop stress contours for the Ecoflex cylinder modeled using an Ogden hyperelastic formulation. (c) Radial and hoop stress contours for the esophageal cylinder modeled using the Marlow hyperelastic formulation. (d) Comparison between theoretical predictions and finite element results for the normalized inner radius as a function of the applied external pressure. (e) Radial stress distribution across the normalized wall thickness showing theoretical and finite element solutions. (f) Hoop (circumferential) stress distribution across the wall thickness comparing theoretical and finite element predictions.

#### Kinematics

Let *R* ∈ [*R*_*i*_ , *R*_*o*_] and *r* = *r*(*R*) ∈ [*r*_*i*_, *r*_*o*_] denote the radial coordinates in the reference and deformed configurations. The principal stretches are

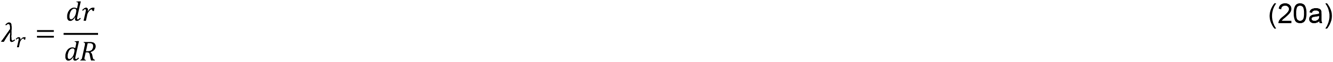

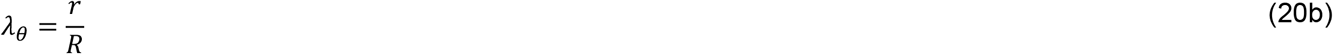

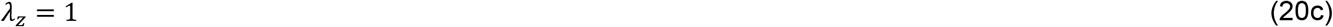

For an incompressible material *λ*_*r*_*λ*_*θ*_*λ*_*z*_ = 1 which gives 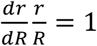. Integration yields

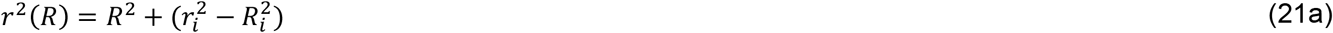

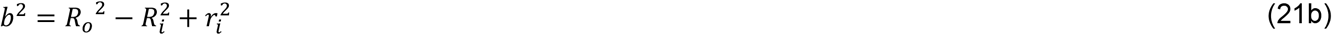

Reducing the problem to a single unknown, *r*_*i*_. Defining 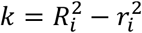 and *R*^2^ = *r*^2^ + *k* the principal stretches become

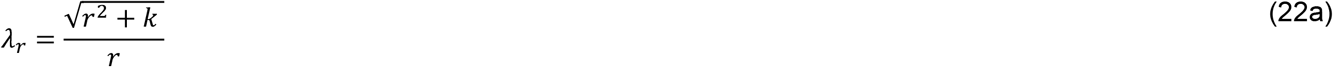

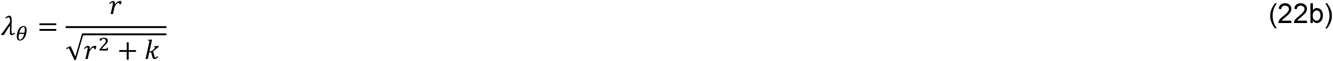

#### Equilibrium

The radial equilibrium equation in the deformed configuration is given by

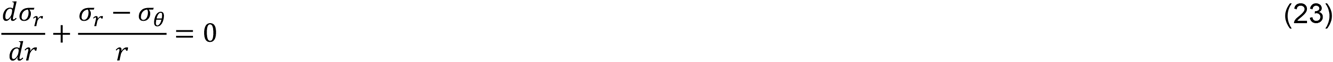

With the boundary conditions given by *σ*_*r*_(*r*_*i*_) = 0 and *σ*_*r*_(*r*_*o*_) = −*P*. Integration from *r* = *r*_*o*_ to any *r* and imposing *σ*_*r*_(*r*_*i*_) = 0 gives the pressure-deformation relation

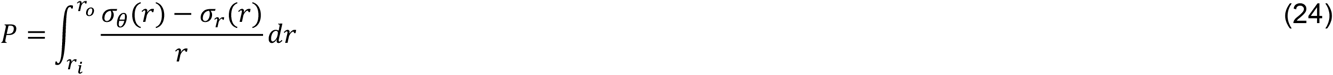

Where *σ*_*θ*_ − *σ*_*r*_ is obtained from the form of the strain energy function.

#### Mooney-Rivlin

For an incompressible Mooney-Rivlin materials, the strain energy function is modified from Section 2.6 as

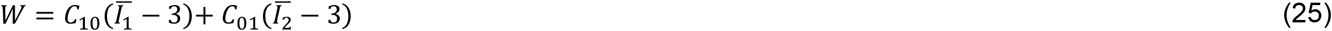

Under plane strain assumptions, *σ*_*θ*_(*r*) − *σr* (*r*) simplifies to

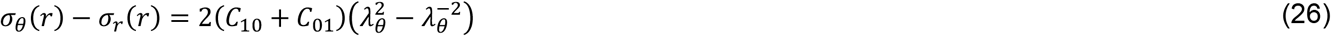

and from the kinematics in Eq. 22b are then expressed as

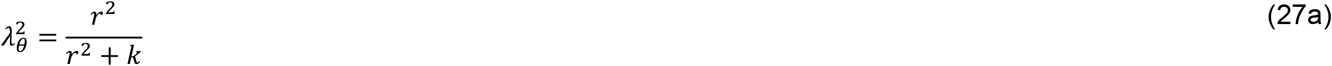

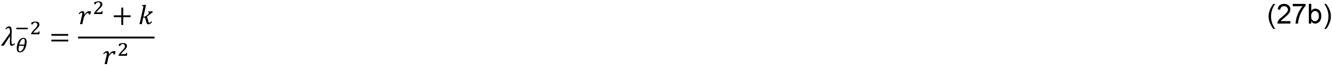

Therefore, the term 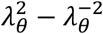 becomes

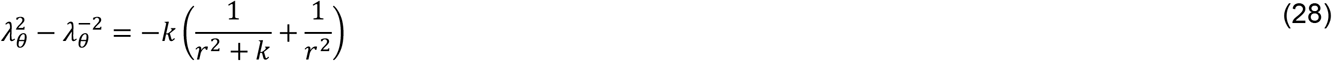

Eq. 26 can be written as

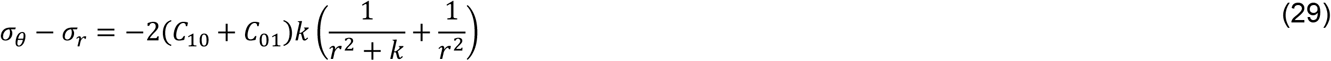

Where *μ* = 2(*C*_10_ + *C*_01_) is the shear modulus. Substituting Eq. 29 into Eq. 24 yields the elementary integral

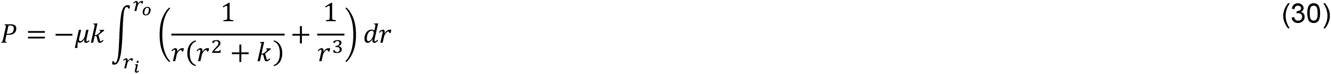

Integrating each team yields

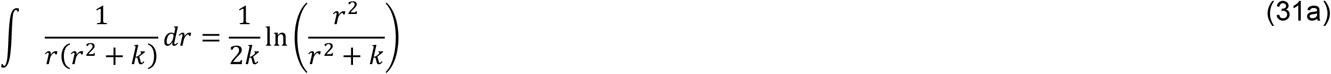

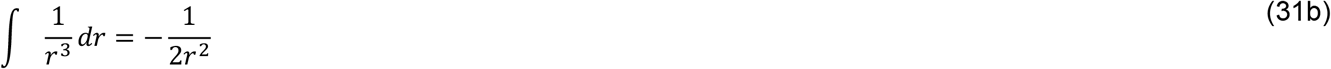

Combining and evaluating the integral yields the implicit equation for the pressure as a function of the deformed radii

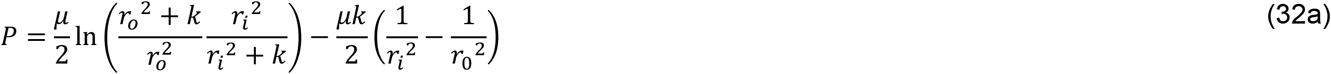

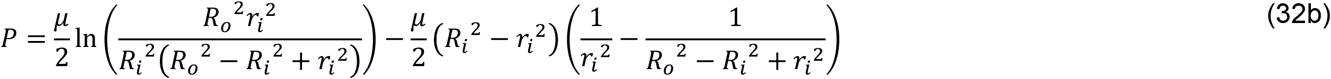

Then, the corresponding radial and hoop stresses for the Mooney-Rivlin material are derived as

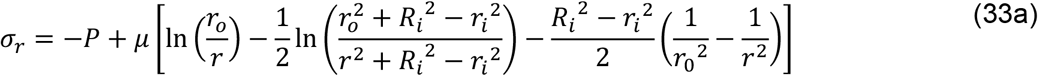

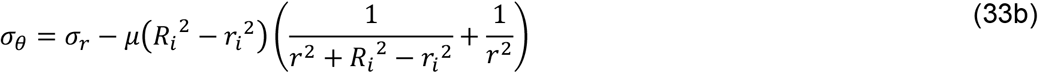

#### Ogden

The Ogden formulation preserves exact finite-deformation kinematics and equilibrium, but the constitutive nonlinearity prevents closed-form evaluation of the pressure integral for general fitted exponents. The problem nevertheless remains semi-analytical, as the full boundary-value problem reduces to a scalar nonlinear root-finding step coupled with one-dimensional quadrature. For an incompressible Odgen material, the strain energy function is modified from Section 2.6 as

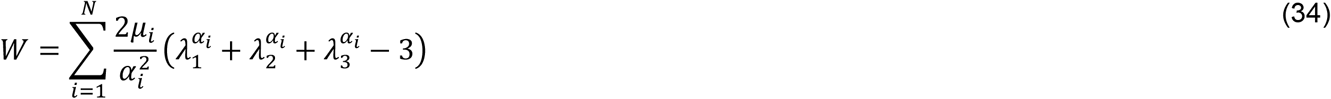

The stress difference (*σ*_*θ*_ − *σ*_*r*_)_Ogden_ is given by

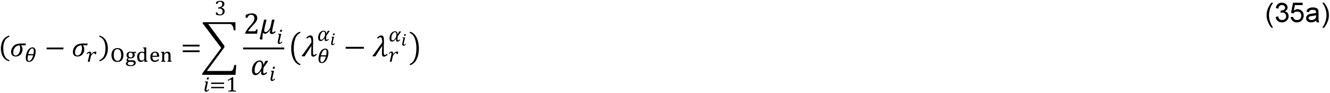

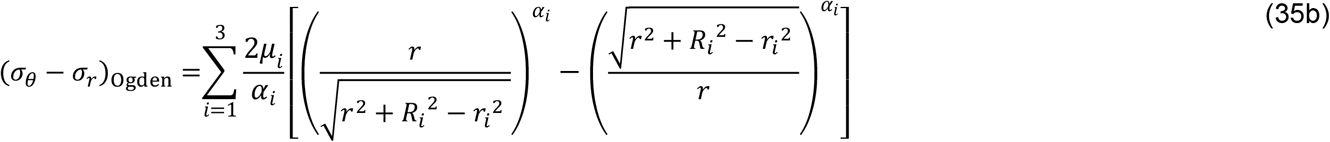

Therefore, the pressure-deformation relations to determine the inner radius *r*_*i*_ becomes

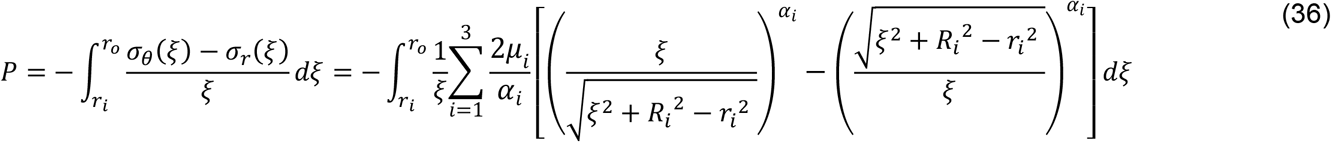

The radial stress at a point *r* is obtained by integrating equilibrium from the inner wall *r*_*i*_ to the point *r* and then the hoop stress is obtained as

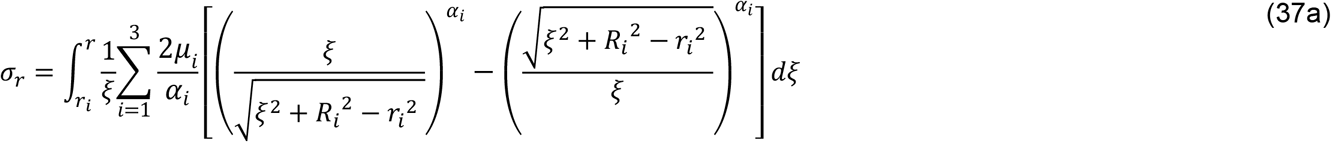

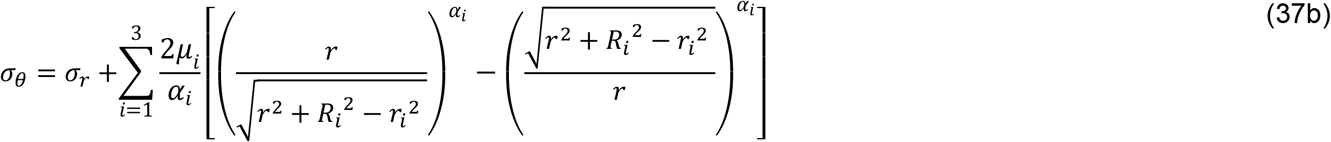

#### Marlow

The Marlow formulation preserves exact finite-deformation kinematics and equilibrium, but the strain energy function *W* = *W*(*I*_1_) is built directly from the uniaxial stress-strain data. For incompressible uniaxial tension the stretches become

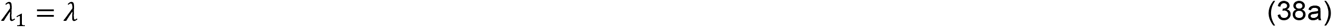

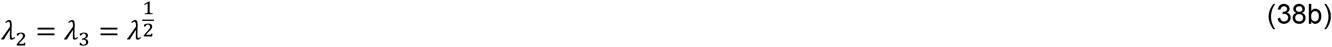

The first invariant *I*_1_ becomes

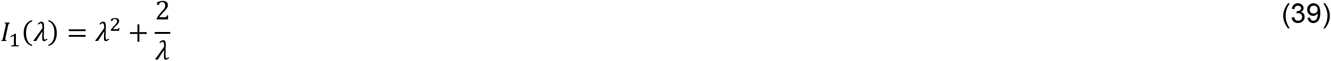

The nominal stress under axial loading is

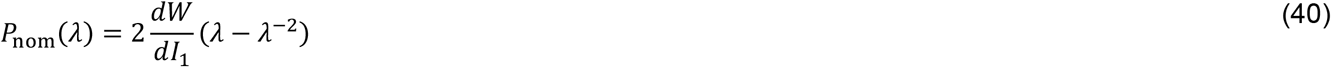

Solving for 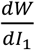 gives the derivative of the strain energy function as

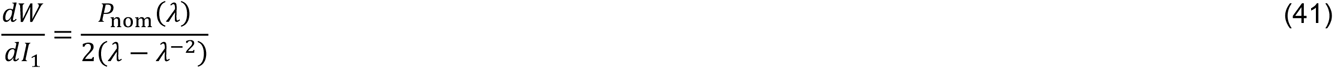

The first invariant of the idealized cylindrical esophagus becomes

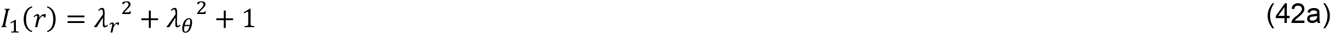

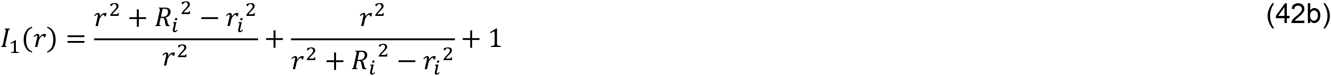

The stress difference (*σ*_*θ*_ − *σ*_*r*_)_Marlow_ is given by

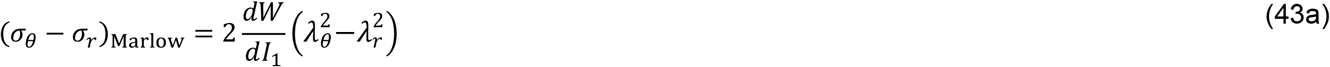

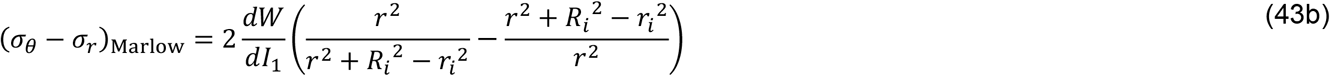

Therefore, the pressure-deformation relationship becomes

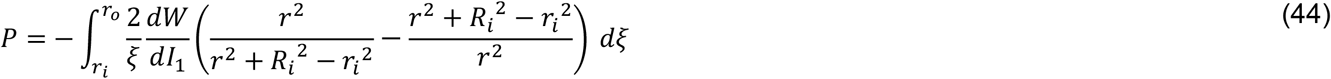

The radial and hoop stresses are obtained as

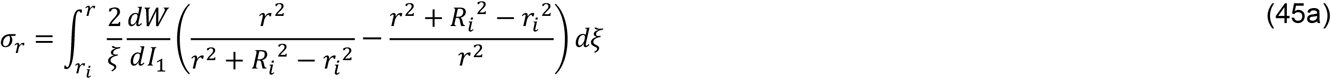

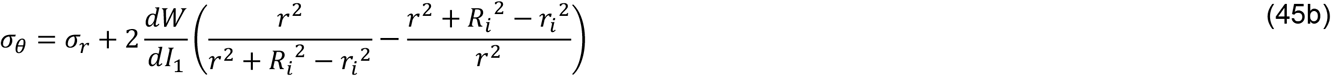

To facilitate practical use, the semi-analytical formulations were implemented in MATLAB, enabling rapid evaluation of the pressure-deformation response across multiple constitutive models. The codes are provided as ready-to-run scripts as part of the supplementary information, allowing non-FEA users (e.g., clinicians) to compute deformation, radial compression, and stress distributions with minimal input data and negligible computational cost. The code can be readily extended to fit additional uniaxial stress–strain datasets for constructing the Marlow strain energy function.

Figure 9a illustrates the axisymmetric deformation kinematics and the corresponding finite element mesh used to validate the semi-analytical solutions. The model is discretized using CPE8H hybrid elements to properly capture the incompressible response. The Ogden and Marlow constitutive models are implemented using the material parameters extracted from Figure 5e-f and Table 1. Figures 9b–c compare the stress distributions predicted for Ecoflex 00-30 and esophageal tissue at external pressures of 5, 10, and 20 kPa. The radial stress *σ*_*r*_ profiles are similar for both materials across the wall thickness, although the esophageal model predicts up to ∼50% higher stresses at larger pressures. In contrast, the hoop stress *σ*_*θ*_ exhibits more pronounced differences, particularly near the inner radius, where the esophageal model predicts stresses up to 2.25 times larger at the lumen. Figure 9d presents a quantitative comparison between the Mooney–Rivlin (Eq. 32b), Ogden (Eq. 36), and Marlow (Eq. 44) formulations, demonstrating excellent agreement between the semi-analytical predictions and the finite element results for the normalized inner radius 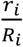. For the Ogden and Mooney–Rivlin models, the decrease in inner radius is nearly linear at small pressures (<5 kPa) before transitioning to a nonlinear response at higher pressures. In contrast, the Marlow model—characterized by the “J-shaped” stress– strain behavior shown in Figure 5f—predicts a significantly stronger contraction, with approximately a 10% reduction at 3 kPa compared to about 2% for the other models. It is important to note that the Marlow model is not extrapolated beyond the available experimental data. Despite this constraint, the model predicts radial compression up to 60 kPa, corresponding to a normalized inner radius of 0.76, compared to 0.62 and 0.64 predicted by the Ogden and Mooney–Rivlin models, respectively. Figures 9e–f present the radial and hoop stress distributions as a function of the normalized radius 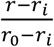 for two representative pressure levels, 5 kPa and 20 kPa. The semi-analytical solutions closely match the finite element predictions shown in Figures 5b–c. The modeling results further highlight the influence of material behavior on the stress response. The esophageal model predicts larger radial stresses that progressively converge toward the Ecoflex response as the normalized radius 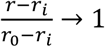. In contrast, the hoop stress distribution differs substantially, with the nonlinear Marlow model producing a pronounced stress decay of approximately 100 kPa across the normalized wall thickness.

## 4. DISCUSSION

This study evaluates how geometric and material modifications to cervical retractor blades influence the mechanical loading imposed on the esophageal wall during anterior cervical discectomy and fusion. A mechanics framework that integrates finite element simulations with controlled experiments quantifies how blade width and compliant interfacial buffering modify the distribution of stress states and contact pressure at the esophageal interface. The results show that relatively simple changes in retractor design alter the local stress state generated during lateral retraction. Increasing blade width and introducing a soft buffer sleeve reduces peak stresses and redistribute loading over a larger contact region. Blade width plays a dominant role in determining the spatial distribution of stress along the esophageal surface. Wider blades increase the contact area between the retractor and the tissue, allowing the imposed retraction displacement to distribute over a larger region. The simulations show that narrow blades produce highly localized loading beneath the blade centerline, whereas wider blades generate smoother stress gradients along the axial direction of the esophageal segment.

The introduction of soft polymeric buffers further modifies the contact mechanics by reducing the stiffness mismatch between the rigid blade and the compliant esophageal wall. In the absence of a buffer, the rigid blade transmits displacement directly to the tissue, producing steep stress gradients and elevated contact pressures. A soft elastomeric layer accommodates part of the imposed displacement through elastic deformation. This additional compliance redistributes the load across the interface and reduces peak stresses transmitted to the esophageal wall to prevent bulging. The predicted stresses remain well below reported ultimate tensile strengths of esophageal tissue, which range from approximately 1.2 to 2.19 MPa under uniaxial loading (Egorov et al., 2002; Vanags et al., 2003). However, the simulated contact stresses approach physiological pressure levels reported in the cervical esophagus and upper esophageal sphincter (6–17 kPa) (Tsou et al., 2021; Cook et al., 1987). Although these loads are unlikely to cause structural failure, localized compression within this range may perturb normal physiological function. Sustained stresses may affect microvascular perfusion within the esophageal wall and contribute to transient dysfunction during postoperative recovery.

Experimental measurements using thin-film pressure sensors support the trends predicted by the simulations. Stepwise retraction experiments show good agreement with finite element predictions for displacements up to approximately 15 mm, indicating that the calibrated hyperelastic material model captures the first-order mechanical response of the phantom. At larger displacements, the simulations predict higher pressures than measured experimentally. This difference likely arises from non-uniform compliance in the sensing system and evolving contact conditions that are not fully captured by the rigid blade idealization. Nevertheless, both experiment and simulation show a monotonic increase in interface loading with increasing retraction displacement. Continuous retraction experiments show that compliant Ecoflex buffer layers reduce the magnitude of the loading signal during retraction, measured as the voltage change between the pre- and post-retraction states. The systematic decrease in voltage signal amplitude with increasing buffer thickness supports the hypothesis that compliant buffering moderates load transfer at the interface by redistributing the imposed displacement over a larger contact region and attenuating localized pressure peaks near the blade edges.

The semi-analytical models provide a mechanistic description of deformation and stress development in the esophageal wall under external compression and enables rapid comparison between constitutive models for soft tissues. The results show that material nonlinearity strongly influences the predicted mechanical response. The Mooney–Rivlin and Ogden models produce similar behavior at low pressures, with an approximately linear reduction in lumen radius followed by nonlinear stiffening. In contrast, the Marlow formulation predicts larger lumen contraction at small pressures. These differences are most pronounced in the hoop stress near the lumen, where the nonlinear tissue response generates higher stresses than those predicted using elastomeric surrogate materials. At higher pressures, however, the radial stress profiles converge, indicating that the influence of constitutive nonlinearity decreases as compression increases. The semi-analytical solutions show excellent agreement with finite element simulations while reducing the problem to a scalar root-finding step and one-dimensional quadrature, enabling rapid evaluation of stress distributions without full numerical simulations.

## 5. LIMITATIONS

Several modeling simplifications were introduced to isolate the effects of blade geometry and interfacial compliance. The 3D and 2D model represent the esophagus as a cylindrical deformable tube with fixed boundary conditions at the proximal and distal ends. This representation neglects surrounding anatomical structures such as the trachea, cervical spine, and connective tissues. These structures may impose additional contact constraints and modify the stress distribution within the esophageal wall during retraction. Consequently, the model may underestimate the multiaxial stress state that develops in vivo. Future models that incorporate multi-organ anatomical assemblies and stress relaxation that could improve the representation of mechanical boundary conditions during surgical retraction. Additional limitations arise from the constitutive representation of the esophageal wall. The present study models the tissue as a hyperelastic material calibrated using phantom measurements. The native esophagus, however, contains multiple structural layers with distinct mechanical functions, including the mucosa, submucosa, and muscularis. These layers contain collagen and muscle fiber architectures that produce anisotropic and viscoelastic mechanical behavior. Prior studies indicate that the collagen-rich submucosal layer contributes significantly to resistance against radial compression and influences transmural stress gradients. Constitutive models that incorporate the non-linearity of fiber-reinforced anisotropy and layered tissue architecture (Yang et.al., 2006 (b)) improve predictions of stress distributions under retraction loading. Equally important is the need to characterize the mechanical response of the esophagus under indentation and retraction loading and to validate the predictions obtained from phenomenological hyperelastic models. Future experiments should characterize transmural stress responses under combined compression and shear, ideally using anatomically realistic ex vivo or in vivo animal models such as porcine cervical esophagus.

The analysis also focuses on the quasi-static mechanical response during displacement-driven retraction and does not consider time-dependent or relaxation effects. In surgical practice, retractors often remain in place for extended time (1-3 hours). Under these conditions, viscoelastic relaxation and creep may alter the stress state within the esophageal wall. Incorporating viscoelastic constitutive behavior would enable evaluation of stress relaxation and strain accumulation during prolonged retraction and provide quantitative guidance on allowable retraction times.

Despite these limitations, the present framework integrates theory, computational modeling, and controlled phantom experiments to quantify the mechanical response of the esophagus during surgical retraction. To the authors’ knowledge, this study represents the first mechanics investigation that combines these approaches to examine esophageal loading during ACDF retraction. The framework establishes a quantitative method for evaluating how surgical design parameters, including blade width and interfacial compliance, influence stress distributions and provides a foundation for assessing potential design modifications to retractor hardware.

## 6. CONCLUSION

This study quantifies the mechanical interaction between cervical retractor blades and the esophageal wall during anterior cervical discectomy and fusion. Finite element simulations show that increasing blade width redistributes contact loads over a larger area and reduces localized stress at the blade-tissue interface. Introducing a compliant polymeric buffer layer further reduces stress concentrations by decreasing the stiffness mismatch between the rigid blade and the compliant tissue. Experimental pressure-voltage measurements using thin-film pressure sensors support these trends and demonstrate reduced loading signals during buffered retraction. Although the predicted stresses remain below reported tissue failure thresholds, their magnitude indicates that localized mechanical loading may contribute to functional impairment in sections of the esophagus following surgery. The combined theoretical, computational, and experimental framework developed here provides a quantitative method to evaluate how surgical hardware design influences stress development during and after retraction. These results motivate and inform mechanics modifications to retractor design that may reduce tissue loading without altering the surgical procedure.

## Declaration of competing interest

The authors declare that they have no known competing financial interests or personal relationships that could have appeared to influence the work reported in this paper.

## Acknowledgements

R.A. acknowledges the seed funding support from the Rice University ENRICH Office and the ASME Haythornthwaite Foundation Research Initiation Grants from the Applied Mechanics Division (AMD). C.L. acknowledges funding support from the Rice Engineering Alumni Student Project Grant Program. The authors thank Alexis (Ella) Lopez for initial discussions on the analytical modeling of esophageal compression and Catherine Stidham for assistance with the experimental setup and experimental data collection.

## Data Availability

All the data is included in the manuscript or in the Supplementary Data.

## Supplementary Information

**Supplemental Figure 1:**
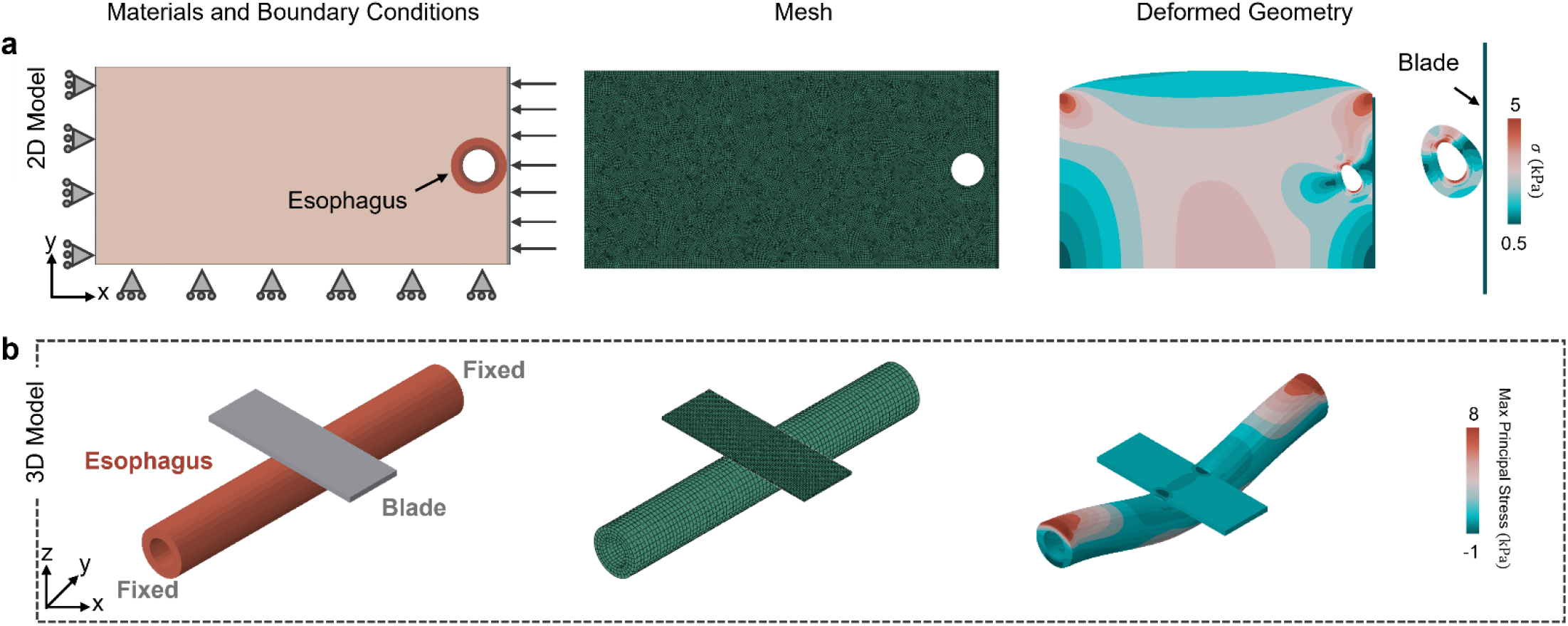
Finite Element Analysis. (a) Two-dimensional finite element model showing the assigned material domains, boundary conditions, computational mesh, and resulting deformed configuration under applied retraction loading. The model captures the cross-sectional mechanical response of the esophagus using hyperelastic material behavior and prescribed blade displacement. (b) Three-dimensional finite element model illustrating the material regions, boundary conditions, discretized mesh, and deformed geometry under simulated surgical retraction. The model extends the 2D formulation to capture the full spatial stress distribution along the esophageal length during blade-induced compression.

## Notes

### Competing Interest Statement

The authors have declared no competing interest.

