## Supplementary Figure 1 for "Mechanics of Esophageal Retraction During Anterior Cervical Discectomy and Fusion"

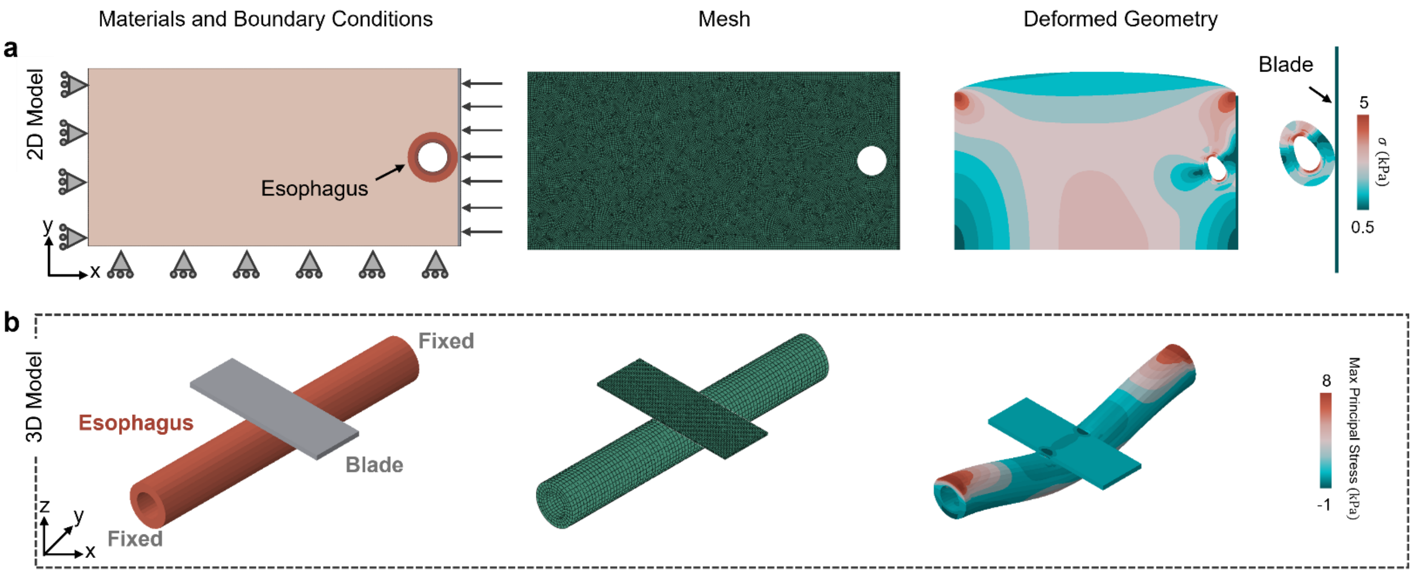


**Supplemental Figure 1 | Finite Element Analysis** (a) Two-dimensional finite element model showing the assigned material domains, boundary conditions, computational mesh, and resulting deformed configuration under applied retraction loading. The model captures the cross-sectional mechanical response of the esophagus using hyperelastic material behavior and prescribed blade displacement. (b) Three-dimensional finite element model illustrating the material regions, boundary conditions, discretized mesh, and deformed geometry under simulated surgical retraction. The model extends the 2D formulation to capture the full spatial stress distribution along the esophageal length during blade-induced compression.
